# Reduced platelet forces underlie impaired hemostasis in mouse models of *MYH9*-related disease

**DOI:** 10.1101/2021.11.10.468045

**Authors:** Juliane Baumann, Laura Sachs, Otto Oliver, Ingmar Schoen, Peter Nestler, Carlo Zaninetti, Martin Kenny, Ruth Kranz, Hendrik von Eysmondt, Johanna Rodriguez, Tilman E. Schäffer, Zoltan Nagy, Andreas Greinacher, Raghavendra Palankar, Markus Bender

## Abstract

*MYH9-related* disease patients with mutations in the contractile protein non-muscle myosin heavy chain IIA display, among others, macrothrombocytopenia and a mild to moderate bleeding tendency. In this study, we used three mouse lines, each with one point mutation in the *Myh9* gene at positions 702, 1424, or 1841, to investigate mechanisms underlying the increased bleeding risk. Agonist-induced activation of *Myh9* mutant platelets was comparable to controls. However, myosin light chain phosphorylation after activation was reduced in mutant platelets, which displayed altered biophysical characteristics and generated lower adhesion, interaction, and traction forces. Treatment with tranexamic acid restored clot retraction and reduced bleeding. We verified our findings from the mutant mice with platelets from patients with the respective mutation. These data suggest that reduced platelet forces lead to an increased bleeding tendency in *MYH9*-related disease patients, and treatment with tranexamic acid can improve the hemostatic function.

**Teaser:** Impaired hemostasis in *Myh9* mutant mice due to reduced platelet forces can be improved by tranexamic acid.

## Introduction

The platelet cytoskeleton ensures normal size and the discoid shape under resting conditions and undergoes rapid rearrangement upon activation. Platelets respond to the biophysical properties of the extracellular environment through integrin-based adhesion sites, which results in actomyosin-mediated contractile forces (*1, 2*). Several inherited platelet bleeding disorders are caused by mutations in key cytoskeletal-regulatory proteins (*3*). *MYH9*-related disease (*MYH9*-RD) is a rare inherited platelet disorder (*4*). The *MYH9* gene encodes the heavy chain of non-muscle myosin 11 A, an actin-binding protein with contractile properties. Heterozygous mutations in the *MYH9* gene in humans lead to macrothrombocytopenia with a moderate bleeding tendency. Depending on the position of the mutations (>30 mutations identified (*5*)), the risk increases for other syndromic manifestations such as renal failure, hearing loss, and pre-senile cataract (*5–7*). Patients with a mutation at amino acid position 702, located in the motor domain of non-muscle myosin IIA, are reported to be most affected for non-hematologic defects and have higher risk for increased bleeding. Mutations located in the rod domain (amino acid positions 1424 and 1841) cause a milder phenotype (*5*). Three different mouse lines with the knock-in mutations Arg702Cys (R702C), Asp1424Asn (D1424N), and Glu1841Lys (E1841K) were generated (*8*), which are the most frequent mutations found in human patients (*9*). These heterozygous point-mutated mice recapitulated key features of human patients, such as macrothrombocytopenia, moderately prolonged bleeding times, decreased ability to retract clots, and non-hematologic defects (*8*). These data show that myosin IIA plays a vital role in platelet production and plug stabilization. However, it is insufficiently understood which factors contribute to the hemostatic defect observed in *MYH9*-RD patients and mutant mice. Given the central role of myosin IIA in force generation and the increased bleeding risk in *MYH9*-RD patients, a better understanding of the underlying biophysical mechanisms in clot formation and its stabilization is warranted (*10*).

Recently, it was demonstrated that biophysical signatures are similar between human and murine platelets (*11*). Therefore, we took advantage of the heterozygous R702C, D1424N, and E1841K point-mutated *Myh9* mice and comprehensively investigated whether altered biomechanical properties might be responsible for the increased bleeding phenotype. While the primary function of mutant mouse platelets was comparable to controls, phosphorylation of the myosin light chain after activation was strongly reduced, the extent of clot retraction decreased, and thrombi were more unstable even when platelet count was adjusted. In line with this, biophysical analyses revealed that *Myh9* mutant platelets generate lower adhesion forces to collagen, lower interaction forces between platelets, and reduced traction forces when spread on fibrinogen-coated micropost arrays. We verified our key findings by analyzing the biophysical function of platelets isolated from *MYH9*-RD patients with the respective mutation. Finally, we observed that treatment with the antifibrinolytic agent, tranexamic acid (TXA), restored clot retraction and reduced bleeding in all three mouse lines. These data suggest that the most common *MYH9-RD* mutations impair the generation of contractile forces by myosin IIA that is necessary to prevent increased bleeding.

## Results

### Unaltered activation, but impaired deformability of *Myh9* mutant platelets

While the role of platelets in hemostatic plug and thrombus formation has been intensively studied from a biological perspective, the mechanobiological aspects are only poorly understood. To investigate the biological and biophysical properties of the contractile protein non-muscle myosin IIA, we capitalized on mice with three different point mutations in the *Myh9* gene. Most results of the D1424N mutation are shown in the main manuscript, results regarding the R702C and E1841K mutations are shown in supplemental results. We confirmed the presence of the protein myosin IIA in platelet lysates from control and heterozygous mutant mice (*Myh9^R702C/+^, Myh9^D1424N/+^, Myh9^E1841K/+^*) by western blot analysis (Supplemental Fig. 1, A and B). The expression levels of D1424N and E1841K mutant nonmuscle myosin IIA are comparable to controls. In contrast, the expression of GFP-tagged human non-muscle myosin IIA with the R702C mutation is 86% of the endogenous expression level, as previously described (*8*) (Supplemental Fig. 1C). *Myh9* mutant mice displayed a significant reduction in platelet count and increased platelet size as determined by a hematology analyzer (Fig. 1, A and B, and Supplemental Fig. 2, A and B; and reference (*8*)). Due to the increased platelet size in mutant mice, control and mutant platelets were gated for a population of similar size in glycoprotein expression and platelet activation studies (Supplemental Fig. 2C). Analysis of expression of prominent platelet surface proteins by flow cytometry revealed moderate differences of some receptors, most notably the GPIb-V-IX complex, even when comparing platelet populations of similar size (Fig. 1C, and Supplemental Fig. 2, C and D). Next, surface exposure of P-selectin and activation of the platelet integrin αIIbβ3 was analyzed after incubation with different agonists. Overall comparable results were obtained for control, and mutant platelets after stimulation of G-protein coupled receptors with ADP, the thromboxane A2 analog U46619, a combination of both, or thrombin (Fig. 1, D and E, and Supplemental Fig. 3, A and B). Similarly, *Myh9* mutant platelets exhibited a comparable degree of activation upon stimulation of the GPVI-ITAM (immunoreceptor tyrosine-based activation motif) pathway with collagen-related peptide (CRP) or convulxin, and of the hemITAM receptor, CLEC2 (C-type lectin-like receptor 2), with rhodocytin. To clarify the role of mutated myosin IIA for platelet shape change and aggregation, *in vitro* aggregation studies were performed. Both agonists, Horm collagen and thrombin, induced a comparable activation-dependent shape change of control and mutant platelets (Supplemental Fig. 4, A and B). Further, *Myh9* mutant platelets showed a normal onset and degree of aggregation (Supplemental Fig. 4, A and B). Next, we investigated whether the mutation in myosin IIA leads to ultrastructural changes in platelets. Transmission electron microscopic (TEM) analysis of mutant platelets revealed a heterogeneous population of platelet size but an otherwise comparable ultrastructure (Fig. 1F, and Supplemental Fig. 4C). Taken together, *Myh9* point-mutated mice display a macrothrombocytopenia, and their platelets can get activated and form aggregates in response to platelet agonists.

**Fig. 1.**
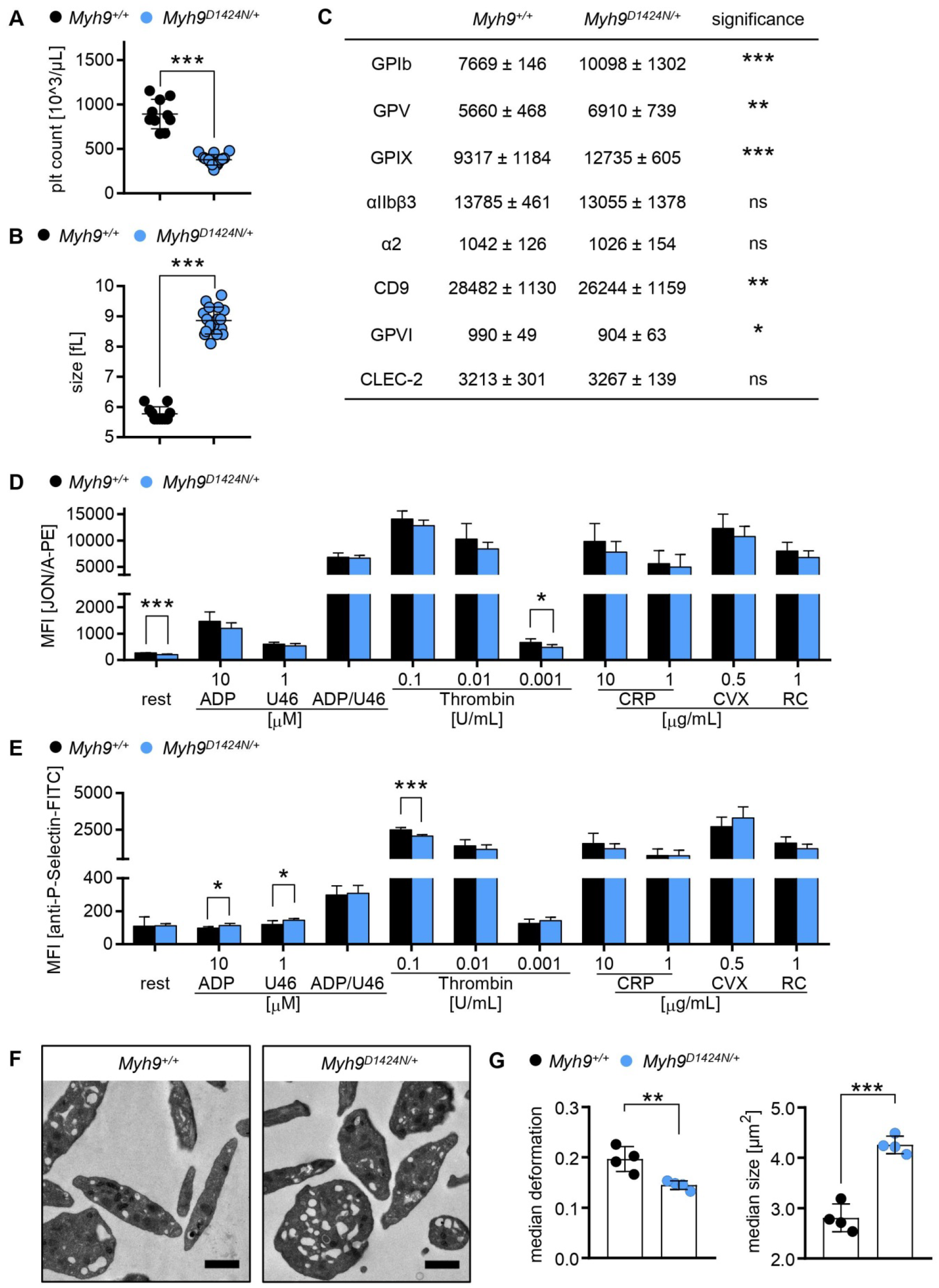
Mutation in *Myh9* gene impairs platelet size and stiffness. (**A**) Platelet count per microliter and (**B**) platelet size of *Myh9^+/+^* and *Myh9^D1424N/+^* mice were determined by a hematology analyzer. (**A** and **B**) Each symbol represents one individual mouse (mean ± S.D.). (**C**) Expression of glycoproteins on the platelet surface was determined by flow cytometry (n=6). (**D**) Activation of platelet αIIbβ3-integrin (JON/A-phycoerythrin [PE]) and (**E**) α-granule release (anti-P-Selectin-fluorescein isothiocyanate [FITC]) were assessed under resting (rest) conditions and upon stimulation with different agonists by flow cytometry (n=6). ADP: adenosine diphosphate; U46: thromboxane A2 analog U46619; CRP: collagen-related peptide; CVX: Convulxin; RC: Rhodocytin. (**C** to **E**) Data are expressed as mean fluorescence intensity [MFI]. (**F**) Transmission electron micrographs from *Myh9^+/+^* and *Myh9^D1424N/+^* platelets; scale bars represent 1 μm. (**G**) Each data point of RT-FDC measurement shows the median deformation or median area of at least 2000 platelets from one subject. Bar plots show mean ± S.D. of platelet deformation or platelet area (n=4).

In contrast, real-time fluorescence deformability cytometry (RT-FDC) measurements revealed a significantly decreased deformation and increased size of *Myh9* mutant platelets (Fig. 1G, and Supplemental Fig. 5, A and B), which points to altered platelet mechanical properties. Since most agonist-induced platelet functions otherwise were normal, we further investigated the structure and function of cytoskeletal components.

### Decreased phosphorylation of the myosin light chain in *Myh9* mutant platelets

To test possible differences in F-actin content, we performed flow cytometry with phalloidin-stabilized resting platelets. Mutant platelets exhibited an increased F-actin content, most likely due to the increased platelet size (Fig. 2A, and Supplemental Fig. 6A). To investigate the organization and rearrangement of the cytoskeleton in *Myh9* mutant platelets, we performed spreading experiments on a fibrinogen-coated surface. Differential interference contrast imaging revealed comparable spreading kinetics of *Myh9^R702C/+^* mutant platelets and slightly faster spreading of *Myh9^D1424N/+^* and *Myh9^E1841K/+^* mutant platelets on a fibrinogen-coated surface (Fig. 2B, and Supplemental Fig. 6B). The increased size might explain the faster adhesion kinetics of these platelets. Similar to control platelets, mutant platelets were able to rearrange the cytoskeleton and form filopodia and lamellipodia, as revealed by platinum replica electron and confocal microscopy (Fig. 2C, and Supplemental Fig. 6, C and D). Phosphorylation of the myosin light chain 2 is responsible for the generation of contractile forces in platelets. Therefore, a capillary-based immunoassay approach was used to detect myosin light chain 2 (MLC2) phosphorylation in resting and thrombin-activated platelets. MLC2 and myosin phosphatase 1 (MYPT1) expression was comparable in control and mutant samples (Fig. 2D, and Supplemental Fig. 1B and 7). However, phosphorylation of MLC2 after activation with thrombin was strongly reduced to 56% in *Myh9^D1424N/+^*, to 40% in *Myh9^R702C/+^,* and to 58% *Myh9^E1841K/+^* mutant platelets compared to *Myh9^+/+^* controls. These data suggest that heterozygous mutations in the motor domain (R702C) and rod domain (D1424N, E1841K) might impair the mutant platelets’ contractile properties.

**Fig. 2.**
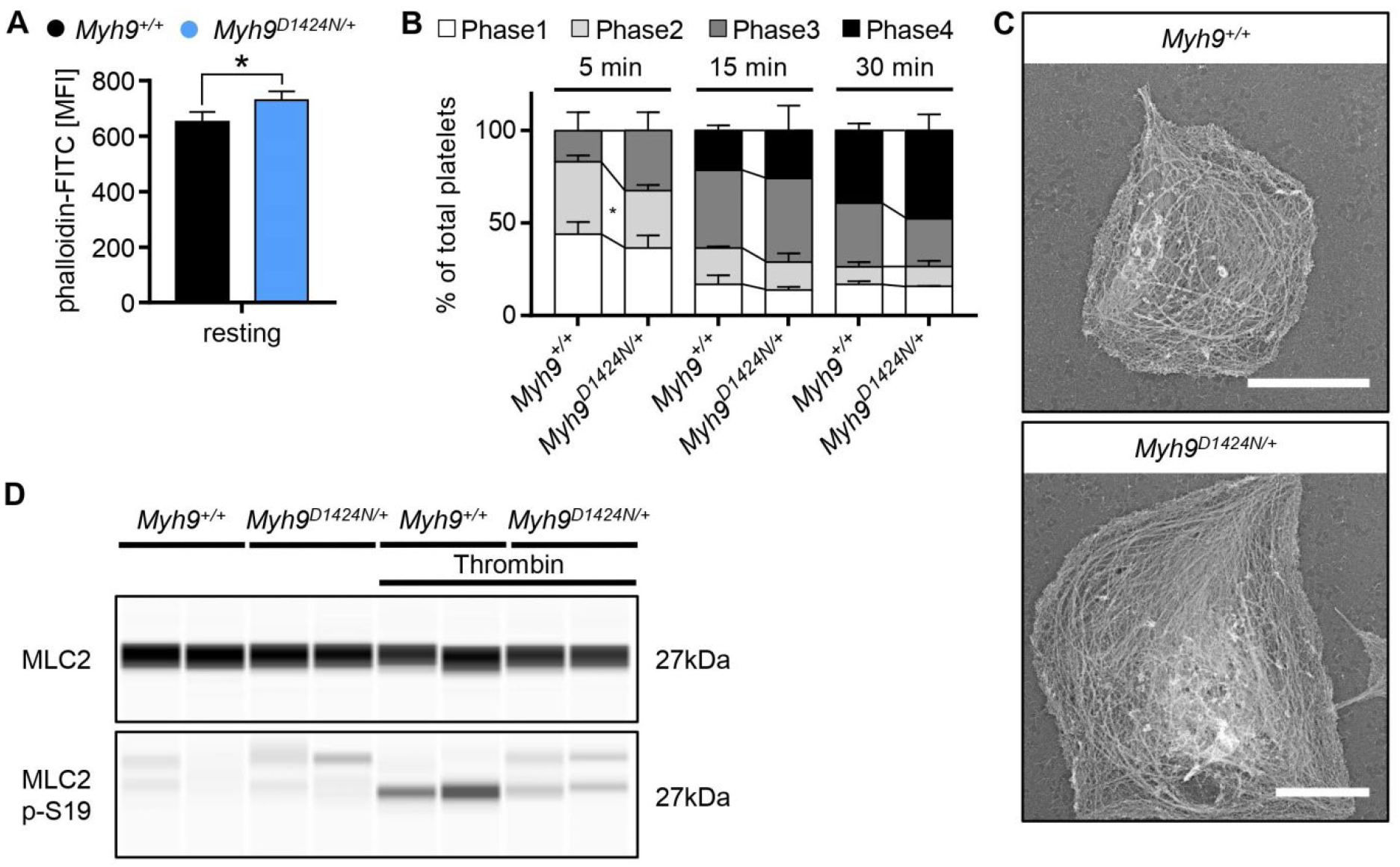
Phosphorylation of MLC2 is decreased in *Myh9^D1424N/+^* platelets. (**A**) F-actin content of resting platelets was measured by flow cytometry after incubation with phalloidin-FITC (n=3-7). The mean fluorescence intensity is shown. (**B**) Statistical analysis of the different spreading phases (phase 1: resting platelets; phase 2: platelets forming filopodia; phase 3: platelets forming lamellipodia and filopodia; phase 4: fully spread platelets) of fixed *Myh9^+/+^* and *Myh9^D1424N/+^* platelets on fibrinogen at different time points expressed as mean ± S.D. (n=2). (**C**) Representative PREM images of the cytoskeleton ultrastructure of platelets spread on fibrinogen in the presence of thrombin (scale bars 2 μm). (**D**) Expression of MLC2 and phosphorylated MLC2p-S19 in resting and thrombin-activated (0.05 U/mL, 1 min) platelets was determined by using an automated quantitative capillary-based immunoassay platform, Jess (ProteinSimple). Representative immunoblot of three independent experiments is shown (n=2).

### Low contractile force generation and impaired clot retraction of *Myh9* mutant platelets

To measure contractile forces, platelets were allowed to adhere and spread on fibrinogen-coated micropost arrays and pulled at posts to different extents. The mean traction force per post beneath each cell was significantly reduced by 38% in *Myh9^D1424N/+^*, 29% in *Myh9^R702C/+^*, and 28% in *Myh9^E1841K/+^* platelets compared to their corresponding controls, as readily perceived from the images of the deformed posts (Fig. 3A, arrows; and Supplemental Fig. 8). However, the total force per cell was not significantly different for *Myh9^D1424N/+^* platelets due to the larger platelet volume of mutant platelets (Fig. 1B), which compensated for the lower reduced intrinsic contractility (Fig. 3A). The total traction forces per single platelet in *Myh9^R702C/+^,* and *Myh9^E1841K/+^* compared to control platelets were significantly lower, indicating that their platelet volume, which is not as increased as for *Myh9^D1424N/+^* platelets (Fig. 1B, and Supplemental Fig. 2B), was not sufficient to compensate for the reduced intrinsic contractility (Supplemental Fig. 8). Platelets spread on a fibrinogen-coated surface were also analyzed for stiffness using scanning ion conductance microscopy (SICM). *Myh9^R702C/+^* and *Myh9^D1424N/+^* platelets displayed a softer appearance on fibrinogen compared to *Myh9^+/+^* platelets, as revealed by a lower Young’s modulus (Fig. 3B, and Supplemental Fig. 9). In contrast *Myh9^E1841K/+^* platelets showed a comparable Young’s modulus to *Myh9^+/+^* platelets (Supplemental Fig. 9). Next, we performed a clot retraction assay since platelet-mediated compaction is an important mechanism to stabilize thrombi (*10*). The extent of clot retraction was significantly different between control and mutant mice. *Myh9* mutant samples showed an impaired clot retraction (*8*), even when the platelet count had been adjusted (Fig. 3C, and Supplemental Fig. 10, A and C). Quantification of the residual clot revealed a heavier, less retracted clot and a corresponding lower volume of residual fluid in *Myh9* mutant samples (Fig. 3D, and Supplemental Fig. 10, B and D). These findings strongly suggest that impaired clot retraction of *Myh9* mutant platelets is due to a defect in the generation of contractile forces and not because of a reduction in platelet count.

**Fig. 3.**
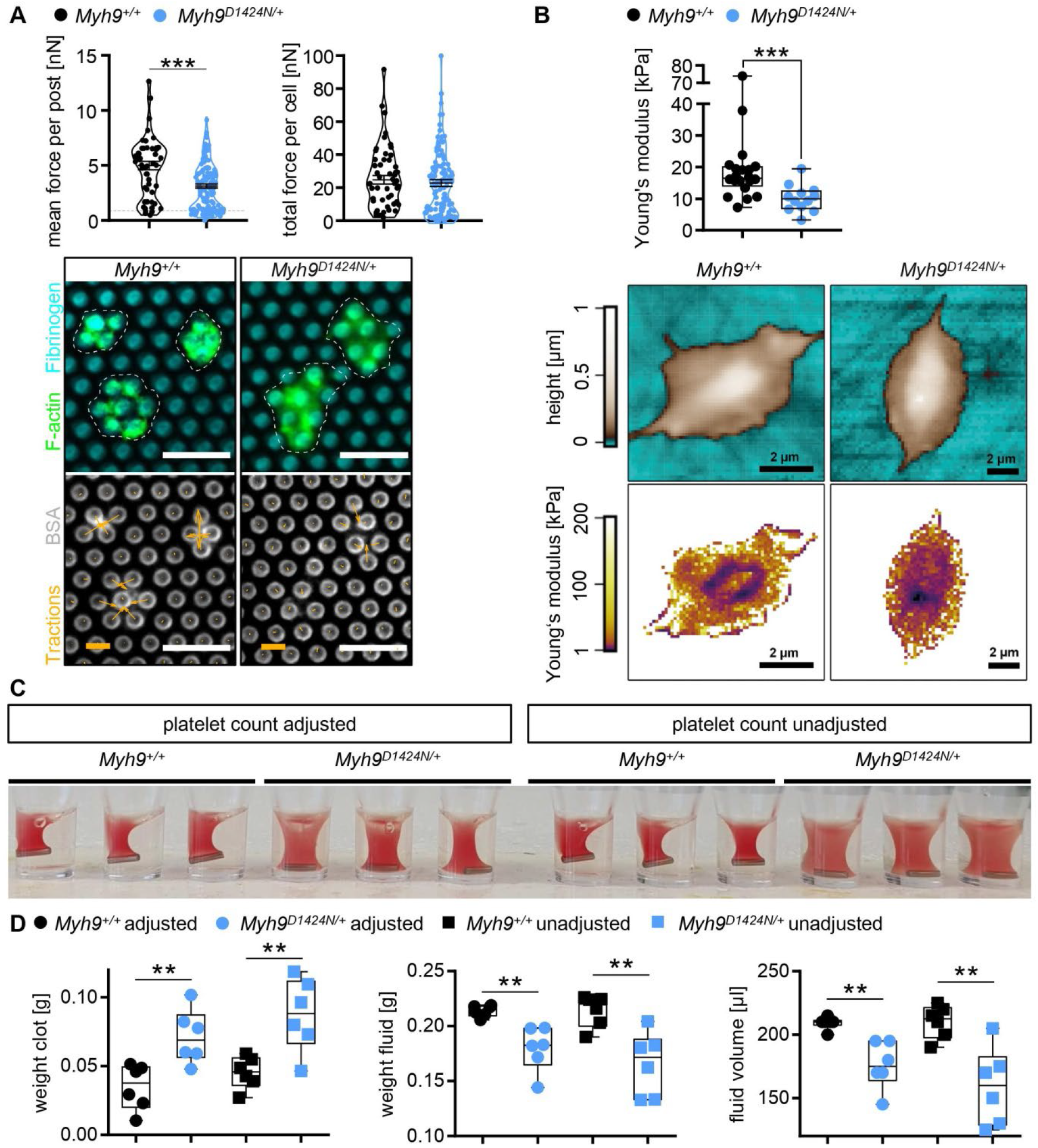
Less contractile forces and reduced clot retraction of *Myh9* mutant platelets. (**A**) Platelet contractile forces were measured per post and the sum of posts per cell (mean ± S.D.; *Myh9^+/+^*: n=51 platelets; *Myh9^D1424N/+^*: n=124 platelets). Representative images of individual platelets (dashed white), stained for F-actin, and its traction on micropost stained for fibrinogen, and BSA (scale bars 5 μm, force bar 10 nN). (**B**) Washed *Myh9^+/+^* and *Myh9^D1424N/+^* platelets spread on fibrinogen in the presence of thrombin were analyzed of their Young’s modulus using scanning ion conductance microscopy. Each symbol represents one platelet (*Myh9^+/+^:* n=20 platelets; *Myh9^D1424N/+^*: n=12 platelets) and box plots represents median ± S.D. Representative images showing the height (upper row) or Young’s modulus (lower row) of *Myh9^++^* and *Myh9^D1424N/+^* platelets. (**C**) Representative image of clot formation at time point 60 minutes (n=3). (**D**) Statistical analysis of weight from residual clot and fluid, and of volume from residual fluid depict median ± S.D. Each symbol represents one individual mouse.

### Lower adhesion and interaction forces of mutant platelets

To investigate the mechanisms of platelet biomechanics under shear flow, we analyzed the effect of the heterozygous point mutations in myosin IIA in an *ex vivo* thrombus formation assay. At first, we studied platelet adhesion to collagen fibers in a whole-blood perfusion assay at shear rates of 1,000 s^-1^. Control and mutant platelets adhered to collagen and formed thrombi both under platelet count adjusted and unadjusted conditions. However, platelets from *Myh9^D1424N/+^* and *Myh9^R702C/+^* mutant mice formed fewer and smaller thrombi than controls (Fig. 4A, and Supplemental Fig. 11A). Surprisingly, platelets from *Myh9^E1841K/+^* mutant mice formed only fewer and smaller thrombi under thrombocytopenic conditions (Supplemental Fig. 11A). Next, platelet adhesion forces on collagen were measured using single platelet force spectroscopy (SPFS). Platelets of *Myh9* mutant mice displayed lower adhesion forces to collagen (Fig. 4B, and Supplemental Fig. 11B; reduced by 69% in *Myh9^D1424N/+^*, by 33% in *Myh9^R702C/+^*, and by 42% in *Myh9^E1841K/+^* platelets). Similarly, interaction forces between two single platelets were reduced for all three mouse lines (Fig. 4C, and Supplemental Fig. 11C; *Myh9^D1424N/+^* by 39%, *Myh9^R702C/+^* by 50%, and *Myh9^E1841K/+^* by 29%). Thrombi formed at a shear rate of 1,000 s^-1^ on collagen were studied for their stiffness by colloidal probe spectroscopy. Thrombi formed by platelets of *Myh9* mutant mice were softer compared to thrombi of control mice as revealed by a decreased Young’s modulus (Fig. 4D, and Supplemental Fig. 11D). In summary, these results suggest that reduced platelet-substrate and platelet-platelet forces lead to reduced thrombus formation under shear.

**Fig. 4.**
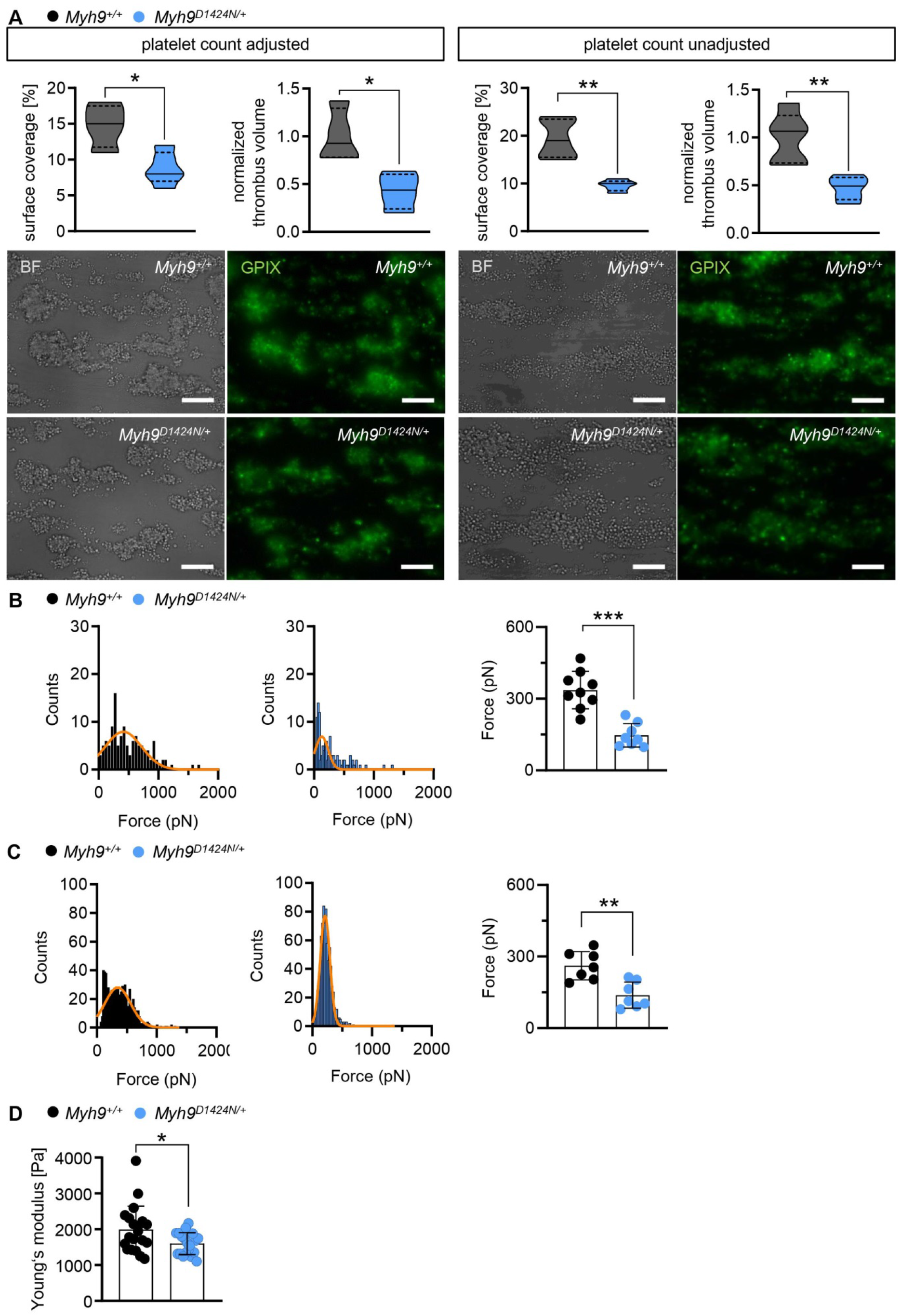
Lower adhesion and interaction forces of *Myh9^D1424N/+^* platelets lead to smaller, and softer thrombi. (**A**) Assessment of platelet adhesion and aggregate formation under flow (1000/s) on collagen of *Myh9^+/+^* and *Myh9^D1424N/+^* samples. Analysis of the surface area covered by platelets (%) and the relative normalized thrombus volume are shown of platelet count adjusted conditions on the left side (mean ± S.D.; n=4) and unadjusted conditions on the right side (mean ± S.D.; n=5). Respective representative images taken at the end of the perfusion time are shown in brightfield and fluorescent images with platelets labeled with the anti-GPIX-antibody (scale bars 30 μm). Single platelet force spectroscopy was performed to determine (**B**) adhesion forces (platelet to collagen) and (**C**) platelet to platelet interaction forces. (**B** and **C**) Representative SPFS curves from one platelet adhering to collagen or interacting with another platelet of *Myh9^+/+^* or *Myh9^D1424N/+^* sample is shown. Each data point of summary graphs (mean ± S.D.) shows one platelet to collagen (n=9) or platelet to platelet (n=7) interaction. (**D**) Each data point of colloidal probe spectroscopy shows the median Young’s modulus of one *Myh9^+/+^* or *Myh9^D1424N/+^* aggregate and bar plots show mean ± S.D. of Young’s modulus (n=4).

### Samples of *MYH9-RD* patients recapitulate the biomechanical phenotype observed in the respective mutant mice

To verify our biophysical results obtained from the analyses of mutant mice, we analyzed deformability and force generation of platelets from two patients with the respective mutations (*MYH9* p.D1424N, *MYH9* p.E1841K). Mechanophenotyping using RT-FDC showed that platelets from patients are stiffer and larger, which is in line with the mouse data (Fig. 5A, and Supplemental Fig. 12A). Next, we assessed platelet adhesion and thrombus formation on collagen under shear. We observed significantly less and smaller thrombi when whole patient blood was perfused over collagen fibers. Moreover, the kinetics of thrombus formation overtime was decreased in the patient sample (Fig. 5B, and Supplemental Fig. 12B). We measured the stiffness of the formed thrombi by colloidal probe spectroscopy. In agreement with the data obtained from *Myh9^D1424N/+^* mutant mice, thrombi were significantly softer for *MYH9* p.D1424N (Fig. 5C). In contrast, stiffness was moderately but not significantly reduced for *MYH9* p.E1841K (Supplemental Fig. 12C). These results show that the mechanical characteristics of human and mouse platelets with point mutations in non-muscle myosin IIA are overall comparable, except for thrombus stiffness of the *MYH9* p.E1841K sample, which shows only a tendency to softer thrombi.

**Fig. 5.**
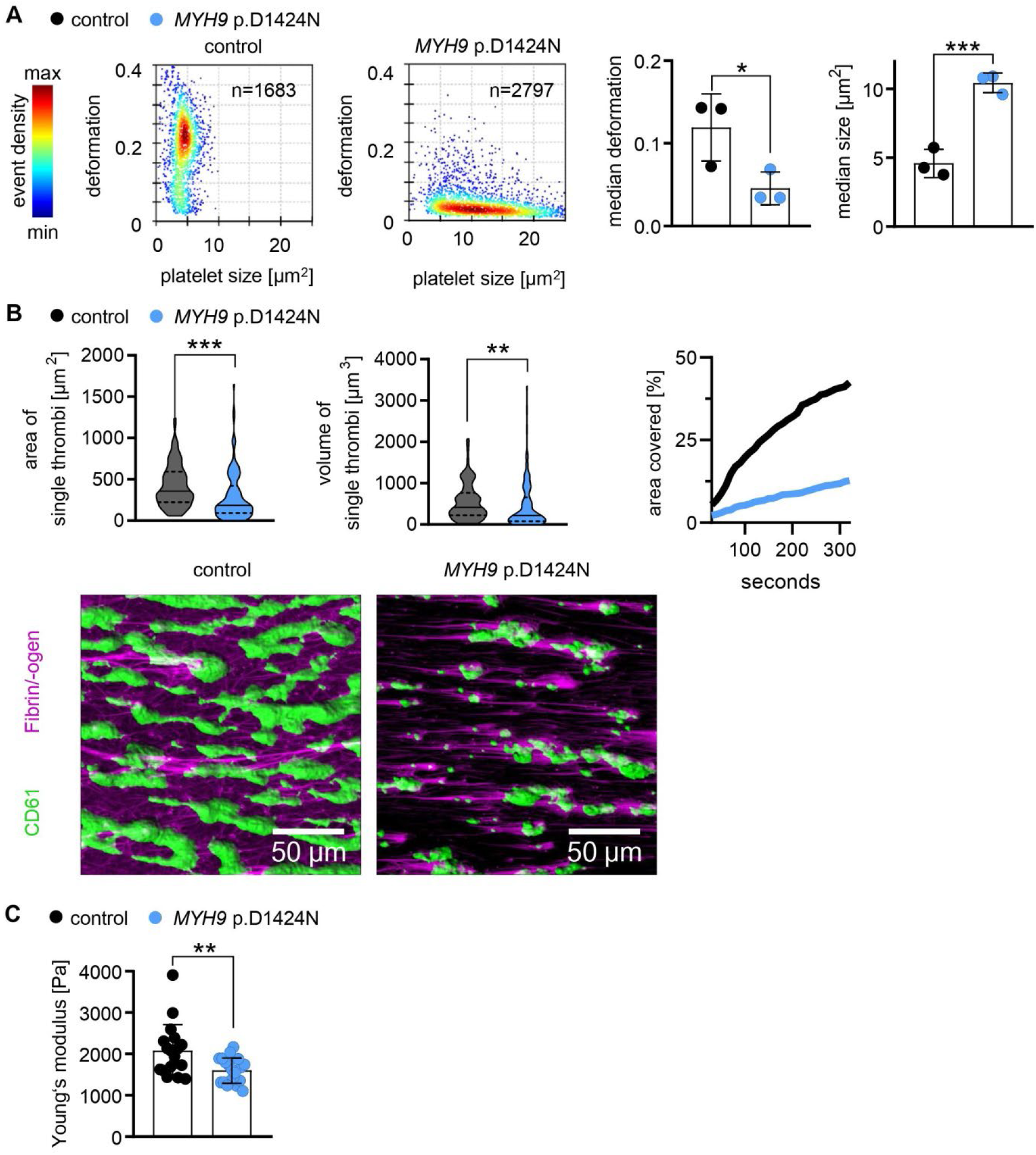
Platelets of patient with the *MYH9* p.D1424N point mutation resemble the biomechanical phenotype of mouse platelets. (**A**) Representative KDE scatter plots from RT-FDC measurements displaying the distribution of single platelet deformation and their corresponding size between single platelets from a healthy individual (control) and from a *MYH9* p.D1424N patient (n= number of single platelets). Summary data points show the median values of individual donors and one *MYH9* p.D1424N patient on three different days and bar plots show mean ± S.D. of platelet deformation and size from non-stimulated platelets. (**B**) Platelet adhesion and aggregate formation under flow (1000/s) on collagen of human patient samples were assessed using a flow chamber. Area of single thrombi and the volume of single thrombi are shown as median ± quartiles of platelet count unadjusted conditions (n=100 single thrombi). Area covered over time shows the mean of thrombi from control and patient blood. Representative images taken at the end of the perfusion time (20 minutes) are shown in fluorescent images with platelets labeled with the anti-CD61-antibody and labeled fibrin/-ogen (scale bars 50 μm). (**C**) Whole blood from healthy individuals and *MYH9* p.D1424N patient was examined by colloidal probe spectroscopy. Each data point shows the median Young’s modulus of one healthy individual or *MYH9* p.D1424N aggregate and bar plots show mean ± S.D. of Young’s modulus.

### Treatment with tranexamic acid improves hemostatic function in mutant mice

*MYH9*-RD patients have an increased bleeding risk. The antifibrinolytic drug, tranexamic acid (TXA), is one option to control bleeding complications in those patients (*5, 6*). We thus hypothesized that gaps of a less compact clot might allow for increased diffusion of plasminogen into the clot, thereby accelerating clot instability and that TXA may help to prevent clot degradation and stabilize the clot of *Myh9* mutant mice. TXA was beneficial in combination with threshold concentration (0.0138 nM) of the recombinant tissue plasminogen activator (rtPA) (Fig. 6A). The addition of TXA in concentrations of 10 mM and 100 μM inhibited the fibrinolytic effect of rtPA and restored clot retraction (Fig. 6A, and Supplemental Fig. 13). As expected (*8*), *Myh9* mutant mice displayed prolonged bleeding times (Fig. 6B, and Supplemental Fig. 14). Strikingly, treatment with tranexamic acid significantly reduced the bleeding time in the three *Myh9* mutant mouse models (Fig. 6B, and Supplemental Fig. 14). In summary, the increased bleeding phenotype due to reduced platelet forces in *Myh9* mutant mice can be compensated by the addition of TXA.

**Fig. 6.**
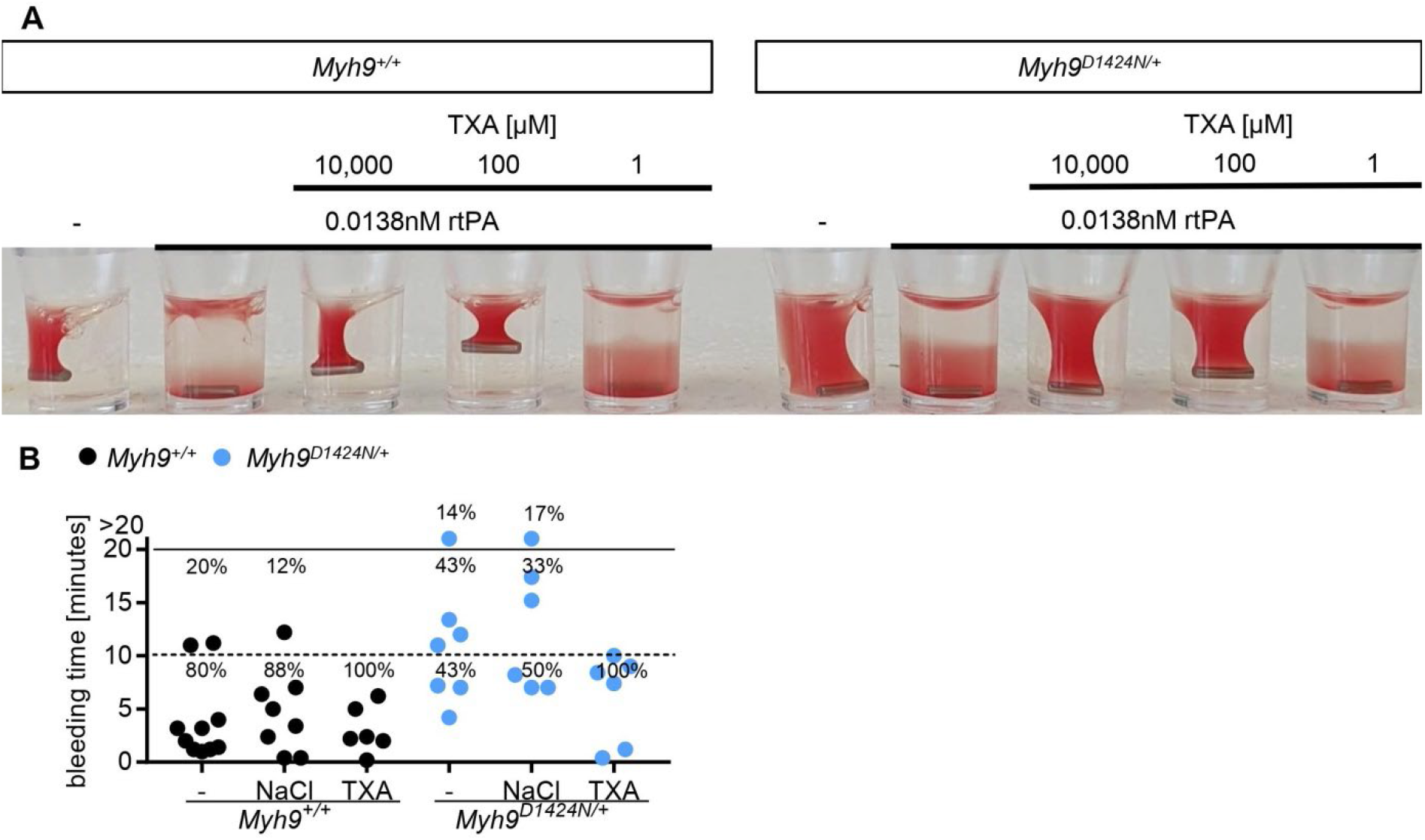
Treatment with tranexamic acid improves hemostatic function in mutant mice. (**A**) Representative image of clot retraction from *Myh9^+/+^* and *Myh9^D1424N/+^* samples treated with rtPA in a threshold concentration where lysis occurs. Addition of TXA in different concentrations 10,000 μM, 100 μM and 1 μM; representative for four independent experiments. (**B**) Tail bleeding times on filter paper of *Myh9^+/+^* and *Myh9^D1424N/+^* mice. Injection of sodium chloride served as injection control. Each symbol represents one individual mouse (mean ± S.D.).

### TXA delays clot lysis of samples from *MYH9*-RD patients

The improved clot stability of mutant mouse samples prompted us to test the effect of TXA on clot formation and stability in patient samples. While the onset of lysis (LOT) measured by thromboelastometry occurred earlier for the human patient sample when adding rtPA, the addition of TXA rescued the defect (Fig. 7A, and Supplemental Fig. 15). Similarly, *ex vivo* thrombus stability of *MYH9*-RD patient platelets under shear was improved in the presence of TXA (Fig. 7, B and C, and Supplemental Fig. 16, A and B). We measured the stiffness of the thrombi with colloidal probe spectroscopy and found that TXA significantly increased the stiffness of thrombi from the *MYH9* p.D1424N sample for an extent of 1.589 (Fig. 7D). Improvement of thrombus stiffness with TXA could be achieved, however, to a lower extent of 1.20 for patient sample with E1841K mutation (Supplemental Fig. 16C). Together, this shows that TXA can also stabilize clots and thrombi of *MYH9*-RD patients.

**Fig. 7.**
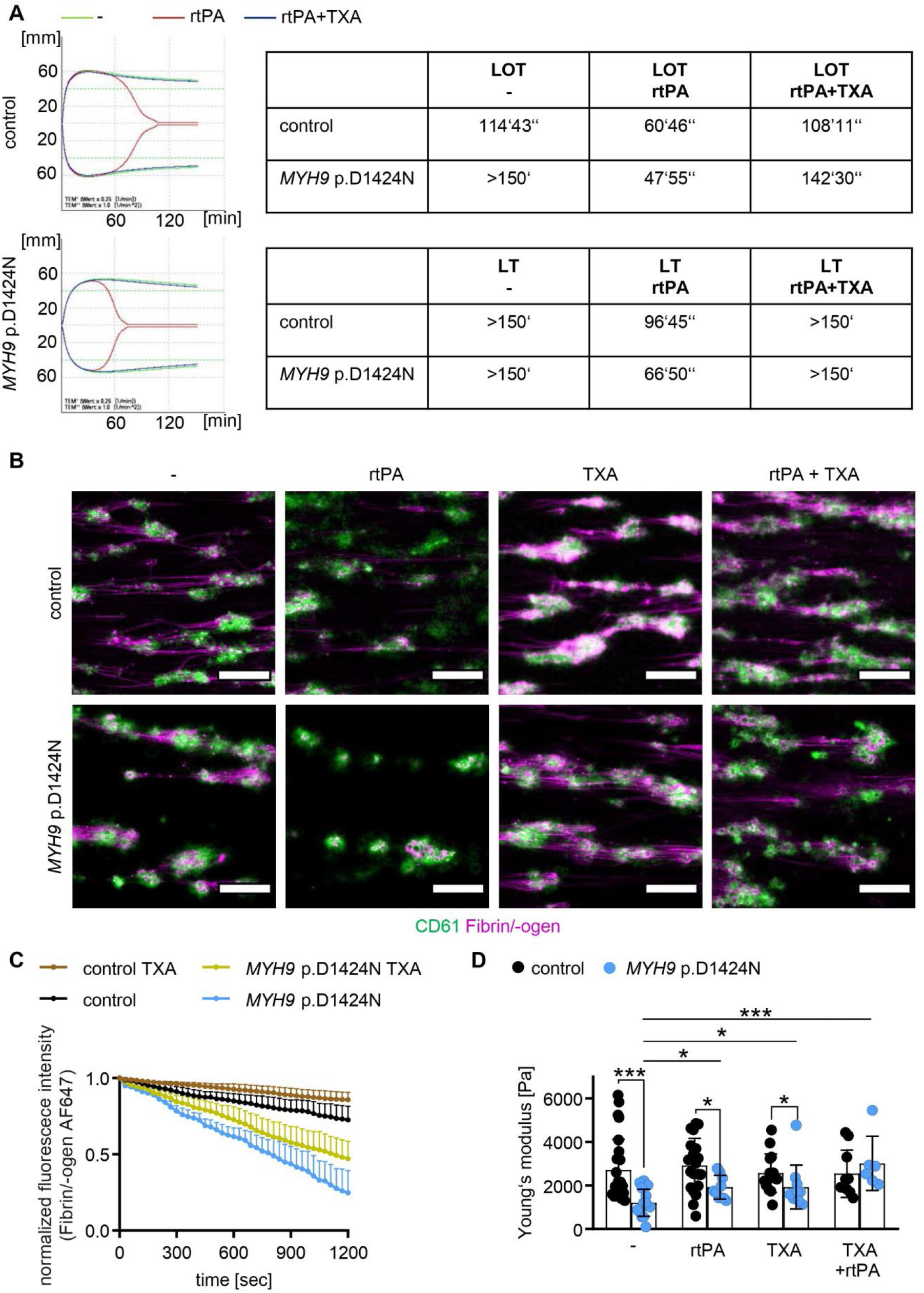
TXA improves clot and thrombus stability of patient sample. (**A**) Lysis onset time (LOT) and lysis time (LT) assessed with ROTEM without treatment (-), stimulation with rtPA, and after addition of TXA to contrast rtPA-induced lysis stimulation. Overlapping of the modified ROTEM analysis curves (green: no treatment; red: rtPA stimulation; blue: rtPA and TXA stimulation). (**B**) Representative fluorescence images of human platelet thrombi (CD61; green) and fibrin (magenta) at 20 minutes after in microfluidic flow chamber at a shear of 100 s^-1^ (scale bars 50 μm). (**C**) Time course of changes in stability of fibrin/-ogen (normalized fluorescence of fibrin/-ogen AF647, mean ± S.D.) on sites of platelet thrombi after addition of TXA (100 μM) compared to non-treated control. (**D**) Whole blood from healthy individuals and *MYH9* p.D1424N patient was examined (untreated, with rtPA, with TXA and with TXA + rtPA) using colloidal probe spectroscopy. Each data point shows the median Young’s modulus of one healthy individual or *MYH9* p.D1424N aggregate and bar plots show mean ± S.D. of Young’s modulus.

## Discussion

While the role of platelets in preventing blood loss is well characterized from a biological perspective, the mechanobiological aspects are only poorly understood. Therefore, we analyzed point-mutated *Myh9* mouse models, which recapitulate clinical manifestations observed in *MYH9*-RD patients. Our study demonstrates that non-muscle myosin IIA-mediated platelet force generation is crucial for sufficient clot compaction and hemostatic function. Our major conclusions are: (1) heterozygous point mutations in the *Myh9* gene lead to reduced platelet adhesion and platelet-platelet interaction forces in mice and *MYH9*-RD patients; (2) impaired clot retraction and prolonged bleeding time are caused by lower platelet force generation, whereas the reduced platelet count is less critical; (3) the mutations R702C, D1424N, and E1841K have a similar impact on the biophysical platelet function, even though the mutation E1841K had less impact on thrombus formation and stiffness; and (4) inhibition of fibrinolysis with TXA can improve the hemostatic function in *Myh9* mutant mice.

We analyzed platelet activation of the *Myh9* R702C, D1424N, and E1841K mutant mice, which was overall comparable to controls when platelets of similar size were compared. The kinetics and extent of platelet aggregation did not differ between mutants and controls. Absent platelet shape change has been observed for myosin IIA-deficient mouse platelets (*12*) and patient platelets (*5, 13*). Surprisingly, we observed initial platelet shape change of the mutant platelets after activation with collagen and thrombin. Myosin IIA-deficient mouse platelets have, in general, a more pronounced phenotype and thus might explain the difference (*12, 14*). However, the difference regarding shape change between the point-mutated mouse platelets and platelets from patients remains unclear. We detected platelet shape change in aggregometry for all three mouse lines and two different agonists, suggesting that the localization of mutation or type of agonist does not explain the difference. Thus far, the point-mutated mouse models recapitulated the key features of *MYH9-RD* patients (reference (*8*) and our findings). However, we cannot exclude that platelets of the patients have a more severe defect in cytoskeletal rearrangement than we observed in the mouse models.

The degree of thrombocytopenia and bleeding varies among *MYH9*-RD patients. It was reported that the patients’ platelet function is normal and severity of bleeding tendency correlates with platelet count (*7, 15*). Interestingly, it was previously demonstrated that the hemostatic function efficiently occurs at unexpectedly low platelet counts when platelets are fully functional, at least in mice (*16*). Therefore, our study addressed the relevance of how strongly mechanical forces contribute to the *MYH9*-related bleeding diathesis. We show that *in vitro* clot retraction of samples from *Myh9* R702C, D1424N, and E1841K point-mutated mice was impaired in both unadjusted and adjusted platelet count conditions, pointing to a less central role of the platelet count. However, clot retraction was still possible to some extent, in contrast to studies using the knockout mice with abrogated clot retraction (*12*). While so far the myosin activity inhibitor, blebbistatin, has been mainly used to analyze platelet force generation (*10, 17–19*), this approach cannot be used to assess the impact of the clinically relevant heterozygote mutations. Therefore we have taken a comprehensive biophysical approach to study the mechanical properties of heterozygously point-mutated *Myh9* platelets. Using SPFS, we demonstrate that *Myh9* R702C, D1424N, and E1841K mutant mouse platelets generate lower adhesion forces on collagen and reduced platelet-platelet interaction forces. We also measured traction forces of mutant mouse platelets on fibrinogen using a micropost deflection assay, which has already been successfully used to measure platelet contractile forces and evaluate platelet functionality (*18–20*). In line with the SPFS results, the average contractile force per post of mutant platelets was decreased. However, the total contractile force generated per platelet of *Myh9^D1424N/+^* mice was comparable to their corresponding controls. One possible explanation might be the enlarged size of *Myh9^D1424N/+^* platelets combined with a higher number of surface receptors. However, for a platelet-rich thrombus, in which contractility per unit volume is the relevant parameter, forces per post must be considered. These findings support the concept (*21*) that the inability of platelets to generate optimal contractile forces is associated with increased bleeding and is the predominant factor over reduced platelet numbers in *MYH9*-RD. Recently, high-throughput hydrogel-based platelet-contraction cytometry, which allows quantifying single-platelet contraction forces, revealed subpopulations of highly contractile human platelets (>30 nN) and that average platelet contractile forces varied considerably amongst healthy donors (*21*). In our study, the average total contractile force generated by wild-type mouse platelets ranged from 15.3 – 26.4 nN. However, a direct comparison of the two approaches cannot be made due to the different experimental conditions.

We further found that point-mutated *Myh9* platelets were less deformable under resting conditions (non-activated, non-spreading platelets) as determined by high-throughput RT-FDC. However, SICM analysis unveiled that spread *Myh9^R702C/+^* and *Myh9^D1424N/+^* platelets are more deformable than controls. While the increased F-actin content might explain the first finding in non-activated mutant platelets, the latter could be due to an altered actomyosin network under the platelet plasma membrane. It is tempting to speculate that the increased deformation of spread mutant platelets might contribute to the reduced traction forces. In support of this, it was shown in mouse embryonic fibroblasts that stiffer cells had a higher net contractile moment and were more prestressed than softer cells (*22*).

We analyzed the spreading behavior of mutant mouse platelets on fibrinogen and found that heterozygous point-mutated *Myh9* mouse platelets form filopodia and lamellipodia to the same extent as controls. This is in contrast to a previous study, which showed reduced lamellipodia formation and observed more unspread D1424N mutant mouse platelets (*23*). We even found slightly increased spreading kinetics of mutant platelets (D1424N, E1841K) at an early time point, which may probably result from the enlarged platelet size and, therefore, earlier adhesion to the activating surface. We recently showed that lamellipodial structures cannot be observed in a blood clot and therefore are dispensable for the hemostatic function and thrombus formation (*24*). Kim *et al.* demonstrated that platelet filopodia extension and retraction are needed for transmitting platelet contractile force and rearranging the fibrin matrix (*25*). In a very recent publication, the cytoskeletal structure in platelet protrusions was analyzed by applying cryo-electron tomography, revealing a nonuniform polarity of actin filaments in protrusions, indicating that this organization may allow the generation of contractile forces (*26*). Our data in this study demonstrate that impaired clot formation is most likely due to reduced mechanical forces and not because of a defect in shape change or cytoskeletal rearrangement. Although we did not directly analyze actin stress fiber formation upon platelet spreading, immunofluorescent and platinum replica electron microscopic images revealed that most mutant platelets were able to form stress fibers on a continuously fibrinogen-coated surface. In contrast, myosin IIA-deficient platelets are unable to form stress fiber-like structures (*12*). This again shows that a complete deficiency of the protein myosin IIA in platelets produces a more severe phenotype than an amino acid change on one position.

In clinics, there are different approaches to treat bleeding complications in *MYH9*-RD patients (*27*). Thrombopoietin receptor agonists are an option for short-term treatment by increasing the platelet count (*28*). Desmopressin can be given to adults before surgery in combination with TXA. In addition, TXA is also used alone to treat excessive bleeding during menses or is locally applied after dental surgery (*6, 29*). Samson and colleagues showed that limited fibrinolysis paradoxically facilitates more efficient clot retraction, and TXA prevents clot shrinkage (*30*). But if TXA prevents clot retraction, why is it beneficial for *MYH9*-RD patients with bleeding complications? We hypothesized that plasminogen could better enter the gaps between platelets and more efficiently lyse the clot due to reduced interaction forces between mutant platelets. TXA, in turn, stabilizes the clot and prevents plasmin-induced clot instability. Indeed, we found that the administration of TXA could reverse the prolonged bleeding phenotype in mutant mice. Moreover, plasmin-induced clot lysis *in vitro* was prevented by TXA at a final concentration of 100 μM, which is expected to inhibit fibrinolysis (*31*). These findings suggest that insufficient hemostatic plug compaction due to reduced forces of *Myh9* mutant platelets can be overcome by TXA-mediated clot stabilization.

In summary, we comprehensively analyzed the molecular and mechanical characteristics of human and mouse platelets with heterozygous point mutations in non-muscle myosin IIA. We found that impaired clot retraction and increased bleeding tendency can be linked to reduced force generation and that interfering with the fibrinolytic system improves hemostatic function.

## Materials and Methods

### Animals

*Myh9* mutant mice (*8*) were purchased from Mutant Mouse Resource & Research Centers (MMRRC). Stock numbers are 036196-UNC (R702C), 036210-UNC (D1424N), and 036698-MU (E1841K). Wild-type littermates were used as controls for the heterozygous mice from each mutant strain because of the different backgrounds. Female and male mice were between 6 and 12 weeks of age. Animal studies were approved by the district government of Lower Franconia, Germany.

### Human blood samples

The use of whole blood and PRP from healthy adult individuals and *MYH9*-RD patients was approved by the ethics committee of the University Medicine Greifswald, Germany. All participants gave written, informed consent.

### Platelet preparation

#### Mouse

Blood was collected in a tube containing heparin (20 U/ml, Ratiopharm) and platelet-rich plasma (PRP) was obtained by centrifugation at 80*g* for 5 min at room temperature (RT). For the preparation of washed platelets, PRP was centrifuged at 640*g* for 5 min at RT. The platelet pellet was resuspended in modified Tyrode’s-HEPES buffer (134 mM NaCl, 0.34 mM Na_2_HPO_4_, 2.9 mM KCl, 12 mM NaHCO_3_, 5 mM HEPES, 1 mM MgCl_2_, 5 mM glucose, and 0.35% bovine serum albumin [BSA; pH 7.4]) in the presence of prostacyclin (0.5 μM) and apyrase (0.02 U/mL). Platelets were finally resuspended in the same buffer without prostacyclin (pH 7.4; 0.02 U/mL apyrase) and incubated at 37°C for 30 min before use.

#### Human

The donors had not taken any medication in the previous ten days before blood collection. Whole blood was collected by venipuncture in BD Vacutainer^®^ Tubes containing acid citrate dextrose solution A (ACD-A) and 3.8% buffered trisodium citrate (Na-Citrate). Whole blood was stored at 37°C for at least 30 min (at an angle of 45° to the horizontal surface), and PRP was transferred to a new polypropylene tube. All experimental measurements were performed within 3 hours of drawing the blood.

### Immunoblotting

Washed platelets were lysed, separated by sodium dodecyl sulfate-polyacrylamide gel electrophoresis and blotted onto polyvinylidene difluoride membranes. Membranes were incubated with the respective antibodies against GAPDH (Sigma) and myosin IIA (Sigma). Horseradish peroxidase-conjugated secondary antibodies and enhanced chemiluminescence solution (MoBiTec) were used for visualization on an Amersham Imager 680 (GE Healthcare) or a Cawomat 2000 IR apparatus (CAWO solutions).

### Immunoblotting with ProteinSimple Jess

Protein levels of MLC2, myosin IIA, MYPT1 and β-actin, and phosphorylation levels of MLC2 were analyzed using an automated capillary-based immunoassay platform (*32*); Jess (ProteinSimple). 5×10^8^ platelets/mL were lysed by addition of equal volume ice-cold 2x lysis buffer (300 mM NaCl, 20 mM TRIS, 2 mM EGTA, 2 mM EDTA, 10 mM NaF, 4 mM Na3VO4, 1% IGEPAL-CA630). The lysis buffer contained 2x Halt Protease and Phosphatase Inhibitor Cocktail, EDTA-Free (Thermo Scientific) to prevent protease and phosphatase activity. Lysates were diluted to the required concentration with 0.1x Sample Buffer 2 (diluted from 10x Sample Buffer 2). Lysates were prepared by the addition of 5x master mix containing 200 mM dithiothreitol (DTT), 5x sample buffer and fluorescent standards (Standard Pack 1) and boiled for 5 minutes at 95°C according to the manufacturer’s instructions. The optimized antibody dilutions and respective lysate concentrations for each antibody (all from CST) are listed below: anti-MLC2 antibody 1:10, 0.1 mg/mL; anti-myosin IIA antibody 1:10, 0.025 mg/mL; anti-MLC2 p-S19 antibody 1:10, 0.4 mg/mL; anti-MYPT1 antibody 1:10, 0.1 mg/mL; anti-β-actin antibody 1:10, 0.1 mg/mL. For most antibodies, either anti-rabbit secondary HRP antibody (DM-001) or anti-mouse secondary HRP antibody was utilized and the chemiluminescent signal was recorded by using the High Dynamic Range (HDR) profile. For detection of myosin IIA, MYPT1 and β-actin signals in the R702C and E1841K knock-in samples, near-infrared (NIR) anti-rabbit secondary NIR Antibody was used with the corresponding fluorescent detection profile. NIR signal is presented in greyscale. All antibodies were diluted in antibody diluent 2. Samples, antibody diluent 2, primary and secondary antibodies, luminol-S and peroxide mix and wash buffer were displaced into 12-230 kDa prefilled microplates (prefilled with Separation Matrix 2, Stacking Matrix 2, Split Running Buffer 2 and Matrix Removal Buffer). The microplate was centrifuged for 5 minutes at 2500 rpm at room temperature. To start the assays, the capillary cartridge was inserted into the cartridge holder and the microplate placed on the plate holder. To operate Jess and analyze results Compass Software for Simple Western was used (version 4.1.0, ProteinSimple). Separation matrix loading time was set to 200 seconds, stacking matrix loading time to 15 seconds, sample loading time to 9 seconds, separation time to 30 minutes, separation voltage to 375 volts, antibody diluent time to 5 minutes, primary antibody incubation time to 90 minutes and secondary antibody incubation time to 30 minutes.

### Transmission electron microscopy

Washed platelets in a concentration of 3 x 10^5^ platelets/μl were fixed with 2.5% glutaraldehyde (Electron Microscopy Science) in cacodylate buffer (pH 7.2, AppliChem). Epon 812 (Electron Microscopy Science) was used to embed platelets. After generation of ultra-thin sections, platelets were stained with 2% uranyl acetate (Electron Microscopy Science) and lead citrate (Electron Microscopy Science). Sections were analyzed on a Zeiss EM900 electron microscope. Platinum replica electron microscopy of spread platelets was performed as previously described (*24*).

### Flow cytometry

Whole blood was withdrawn from anaesthetized mice into heparin and diluted in Tyrode’s-HEPES buffer. To determine glycoprotein expression blood was incubated for 15 minutes with respective fluorophore-conjugated antibodies. For activation studies, washed blood was resuspended in calcified Tyrode’s-HEPES buffer (2 mM Ca^2+^) after washing twice with Tyrode’s-HEPES buffer. Platelets were incubated with agonists for 15 min and stained with fluorophore-labeled antibodies for 15 min at room temperature. For F-actin content analysis, washed platelets were incubated with an anti-GPIX antibody derivate labeled with Dylight-649 (20 μg/ml, Emfret). Cells were fixed in 10% paraformaldehyde (PFA), centrifuged and resuspended in Tyrode’s-HEPES buffer in the presence of Ca^2+^ and 0.1% Triton X-100. Permeabilized cells were stained with 10 μM phalloidin-fluorescein isothiocyanate (FITC) for 30 minutes. Measurements were performed on a FACSCalibur (F-actin content of mouse line D1424N) or FACSCelesta (all others)(BioSciences, Heidelberg, Germany).

### Aggregometry

Washed platelets (160 μL with 0.5×10^6^ platelets/μL) were analyzed in the presence (collagen) or absence (thrombin) of 70 μg/mL human fibrinogen (Sigma). Light transmission was recorded on a four-channel aggregometer (Fibrintimer, APACT, Hamburg, Germany) for 10 min and expressed in arbitrary units, with buffer representing a light transmission of 100%.

### Platelet spreading

Coverslips were coated with 100 μg/ml human fibrinogen (Sigma) for 2 hours at 37°C. After blocking with 1% BSA in PBS coverslips were washed thrice with PBS. Platelets (3 x 10^5^ platelets/μl) were allowed to spread on the coated surface after the addition of 0.01 U/ml thrombin (Roche) and Ca^2+^. After 5, 15 or 30 minutes, platelets were fixed with 4% PFA and permeabilized with IGEPAL CA-630. Images were taken with a Zeiss incubation microscope (100x objective, DIC).

### Immunostaining of platelets

Fixed and permeabilized platelets were stained for myosin IIA using anti-myosin IIA (M8064 Sigma; 1:200 in PBS) and donkey anti-rabbit IgG-Alexa 546 antibodies (1:350 in PBS). F-Actin was stained using phalloidin-Atto647N (A22287 Invitrogen; 1:500 in PBS) and α-tubulin using Alexa488-conjugated anti-α-tubulin antibodies (sc-23948 Santa Cruz; 1:500 in PBS), respectively. Visualization was performed using a Leica TCS SP8 confocal microscope (100x oil objective; NA 1.4). Confocal microscopy was performed with HC PL APO CS2 objectives. Depending on the stainings, a set of monochromatic lasers (488 nm; 561 nm; HeNe 633 nm) tuned to specific wavelengths were used. Detection filters (PMT Trans or ultrasensitive Hybrid detectors) were set to match the spectral properties of fluorochromes.

### Clot retraction

PRP was filled up to a volume of 250 μl with Tyrode’s-HEPES buffer to reach a platelet concentration of 3 x 10^5^ platelets/μl. To investigate clot retraction with unadjusted platelet count 50 μl PRP was filled up to 250 μl with Tyrode’s-HEPES buffer and 1 μl of red blood cells. After the addition of 0.2 U/ml thrombin (Roche) and 20 mmol/l CaCl_2_ clot retraction was observed over a time period of 60 minutes. To investigate treatment with rtPA and TXA, concentrations of 100 μM and 10,000 μM of TXA (Merck) and 0.0138 nM rtPA (Abcam) were used and added together with thrombin and CaCl_2_.

### Bleeding time

Mice were anesthetized and a 1 mm segment of the tail tip was removed with a scalpel. Tail bleeding was monitored by gently absorbing blood with filter paper at 20-second intervals without making contact with the wound site. When no blood was observed on the paper, bleeding was determined to have ceased. Otherwise, experiments were stopped after 20 minutes. Bleeding times exceeding 20 minutes were excluded in the statistical analysis. TXA (Merck, 10 μg/g) or sodium chloride, as control, was injected intravenously 5 minutes before cutting the tail for bleeding time experiment.

### Platelet adhesion under flow

#### Mouse

Heparinized whole blood was diluted in a ratio of 2:1 in Tyrode’s-HEPES buffer containing Ca^2+^. Whole blood or mixed platelet count adjusted blood (based on platelet number) was incubated with 0.2 μg/ml of an anti-GPIX antibody derivate conjugated with Dylight-488 at 37°C for 5 minutes. Coverslips were coated with 200 μg/ml Horm collagen (Nycomed) at 37°C over night and blocked with 1% BSA in PBS for 30 minutes. Whole blood was perfused over cover slips in the flow chamber (slit depth 50 μm) at shear stress equivalent to a wall shear rate of 1000 s^-1^. Videos and images were taken with a Zeiss Axiovert 200 inverted microscope (63x objective). Analysis was performed using ImageJ software.

#### Human

*Ex vivo* thrombus formation assays was performed at a wall shear rate of 1000 s^-1^ on collagen-passivated surfaces (200 μg/mL HORM collagen type I from horse tendon; Nycomed) in a microfluidic parallel platelet flow chamber (on μ-Slide VI 0.1 with physical dimensions: 1 mm width, 100 μm height, and 17 mm length Ibidi GmbH, Germany). To visualize thrombus formation, prior to perfusion, platelets in ACD-A anticoagulated whole blood were stained with FITC-labelled anti-human CD61 antibody (Clone: RUU-PL7F12, Cat No: 340715 BD Pharmingen, USA, used at final ratio 1:100). Fibrin formation was visualized by spiking whole blood with human fibrinogen conjugated to Alexa Fluor 647 (Cat. No. F35200, Invitrogen, used at final concentration of 7.5 μg/mL). Whole blood was recalcified immediately before perfusion. Time-lapse confocal imaging (intervals of 10 seconds per image) was perfomed on a Leica SP5 confocal laser scanning microscope (Leica, Wetzlar, Germany) equipped with HCX PL APO λ blue 40.0×/1.25 oil objective. For image acquisition, fluorophores (FITC and Alexa Fluor 647 were excited with Argon-Krypton (488 nm and Helium-Neon (HeNe, 633 nm) laser lines, respectively, that were selected with an acoustooptic tunable filter (AOTF). Fluorescence emission was collected between 505-515 nm for FITC (detector HyD), and 640-655 nm for Alexa Fluor 647 (detector HyD). To assess the impact of TXA in some experiments, whole blood was preincubated with TXA (100 μM) for 10 minutes prior to perfusion over collagen. Impact of rtPA (137 ng/ml) on fibrin degradation was assessed under shear flow of 100 s^-1^, added after 10 minutes of thrombus formation. Quantitative assessment of platelet adhesion and thrombus formation was performed to obtain the percentage area covered by thrombi over time by computational image analysis using the surfaces creation wizard algorithm in Bitplane Imaris version 7.65 (Oxford Instruments, Abingdon, United Kingdom). Experiments were performed according to International Society on Thrombosis and Haemostasis Scientific and Standardization Committee (ISTH SSC) subcommittee on Biorheology recommendations (*33*).

### Real-time fluorescence deformability cytometry (RT-FDC)

#### Mouse

25 μL of whole blood bled in citrate was suspended in 425 μL CellCarrier B. The measurement was stopped after achieving 10000 single platelet count (hard gate Area 0 – 40 μm^2^). Using the Shape-Out analysis software (https://github.com/ZELLMECHANIK-DRESDEN/ShapeOut2/releases/tag/2.3.0 Version 2.3, Zellmechanik Dresden, Germany), kernel density estimation (KDE) plots of event density were generated, and statistical analysis using Holm-Sidak method was performed to determine the median values for platelet deformation and their size. The range area ratio was limited to 0 – 1.1 and the cell size to 0 – 10 μm^2^ for the analysis.

#### Human

Platelets in PRP were labeled with a mouse anti-human monoclonal antibody CD61-PE (Beckman Coulter). Incubation was performed at room temperature for 10 minutes in the dark. Deformation measurements were performed in a microfluidic chip with a constriction of 15 μm x 15 μm cross-section and a length of 300 μm (Flic15, Zellmechanik Dresden, Germany). RT-FDC measurements were carried out in buffer CellCarrier B (Zellmechanik Dresden, Germany), which is composed of 0.6% (w/v) methylcellulose in PBS (without Ca^2+^, Mg^2+^). Here, 50 μL of immunofluorescently labeled human PRP was suspended in 450 μL CellCarrier B. The human PRP suspension was then driven through the microfluidic chip at flow rates of 0.006 μl/s, and the measurement was stopped after achieving 5000 single platelet count (hard-gate 150-33000 arbitrary units, A.U. for CD61-PE of fluorescence intensity) or after 10 min. RT-FDC data was acquired using the ShapeIn software (Version 2.0, Zellmechanik Dresden, Germany). Using the Shape-Out analysis software, kernel density estimation (KDE) plots of event density were generated, and statistical analysis was performed to determine the median values for platelet deformation and their size. The range area ratio was limited to 0 – 1.1 and the cell size to 0 – 25 μm^2^ for the analysis.

The RT-FDC setup (AcCellerator, Zellmechanik Dresden, Germany) is built around an inverted microscope (Axio Observer A1, Carl Zeiss AG, Germany) mounted with a Zeiss A-Plan 100x NA 0.8 objective (*34, 35*).

### Single platelet force spectroscopy (SPFS)

Silicon CSC12, 0.6 N/m tipless cantilevers (MicroMasch, Tallin, Estonia) and Glass-bottom 35 mm dishes (I BI DI, Martinsried, Germany) were exposed to UV cleaner (Bioforce Labs, Ames, IA, USA) for 30 min. Before coating, the cantilever spring constants were independently measured by a thermal tune procedure (JPK). Cantilevers were incubated with 50 μg/ml collagen G for 3h at 37°C and rinsed three times with Tyrode’s buffer. An aliquot of 15 x 10^3^ platelets/μl in Tyrode’s buffer containing 1.0 mM CaCl_2_ and 0.5 mM MgCl_2_ was dropped onto the passivated glass right before each measurement (10 min, RT). Unbound platelets were removed by Tyrode’s buffer. To immobilize a single platelet on the cantilever, the collagen passivated-cantilever was brought into contact (setpoint 200 pN) with a non-activated platelet on BSA passivated glass slide. It was waited until a single platelet firmly adhered to the ventral surface at the tip of the cantilever. The cantilever with the adhered platelet was then moved to HORM collagen passivated surface and single platelets firmly attached to the HORM collagen for platelet-substrate and platelet-platelet interaction SPFS measurements, respectively. All measurements were carried out in Tyrode’s with Ca^2+^, glucose and BSA using a JPK NanoWizard 3 (JPK, Berlin, Germany). Force distance (F-D) curves were recorded with a Z-length of 7 μm and a setpoint value of 200 pN to control the maximal force of the cantilever against the surface. A velocity of 15 μm/s was used for all measurements to avoid merging of two platelets during contact and to rupture completely two platelets from each other. For each passivated-substrate, the last 500 force-distance curves were taken from 7 to 10 single platelets.

### Colloidal probe spectroscopy

To determine the Young’s modulus of platelet aggregates, indentation experiments were performed by means of colloidal force spectroscopy using an atomic force microscope (Nanowizard 3, JPK Instruments). The atomic force microscope is combined with an optical system comprised of an inverted optical microscope (IX8, Olympus; 20x objective). Gold-coated cantilevers (Cat. No. CP-qp-CONT-AU-A-5, NANOANDMORE GmbH, AFM tip: sphere (1.5 μm – 3 μm)) coated with 8.45 mM poly(ethylene glycol)methyl ether thiol (Mn 800, Cat. No. 729108, Sigma Aldrich) solution for 2 h at room temperature were used. Measurements were carried out in suspension buffer (136 mM NaCl, 0.41 mM Na_2_HPO_4_, 2.65 mM KCl, 12 mM NaHCO_3_, 2.13 mM MgCl_2_, 2.0 mM CaCl_2_, 5.5 mM D-glucose, and 0.35% BSA, pH 7.4). Before each experiment the sensitivity and spring constant of each cantilever were determined individually in suspension buffer. For sensitivity calibration the cantilever deflection upon contact with a freshly cleaved mica surface was analyzed (ensemble average 23,3 ± 3.1 nm/V) and spring constant was measured by thermal noise technique (ensamble average 98 ± 30 mN/m). 1 ml ACD-A human whole blood was treated with 7.5 μl 1 M CaCl_2_ x 2 H2O and 1.88 μl 2 M MgCl_2_ x 6 H2O solution. Heparinized whole blood was used for the mouse experiments. Coverslips, which were coated with 80 μg/ml Horm collagen (Nycomed) at 37°C overnight and blocked with 1% BSA in PBS for 10 minutes, were installed into a flow chamber with a slit depth of 55 μm (Cat. No. 20000209, straight channel chip, microfluidic ChipShop). Recalcified whole blood was perfused at shear stress 1000 s^-1^ for 5 minutes. Coverslips were washed three times with suspension solution. The cantilever was brought into contact with platelet aggregates at a speed of 2.7 μm s^-1^ with a force setpoint defined at 2 nN. Measurements were performed at room temperature in suspension buffer. Platelet Young’s modulus *E* was calculated from AFM indentation force curves by applying the Hertz model which describes how the force increases as the AFM probe pushes into the cell. In short, cantilver deflection force *F* was fitted to the equation 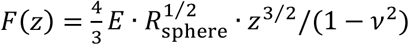. Here, *R*_sphere_ is the radius of the AFM sphere probing the cells, *z* is the cell indentation and the Poisson’s ratio *ν* was set to 0.5 assuming cell volume conservation. Details can be found elsewhere (*36, 37*). Five platelet aggregates with at least 25 force curves per aggregate were recorded per coverslide and each force curve was analyzed individually. Thus, for each experimental condition platelet Young’s modulus is given as the mean value of more than 100 individual force curves.

For rtPA measurements, 500 μl of 137 ng/ml rtPA (Cat No. ab92637; Abcam) was added to the coverslide immediately after measurement of the untreated or TXA sample and incubated for 5 minutes. For TXA measurements, 1 ml whole blood was incubated with 100 μM TXA (Merck) for 10 minutes and further treated as described above.

### Micropost assay

Micropost arrays were fabricated from hPDMS (#PP2-RG07; Gelest Inc) by replica molding from negative molds. Micropost arrays contained cylindrical posts with 1.05 μm diameter and 2.6 μm height arranged on a hexagonal grid with 2.0 μm center-to-center spacing. The Young’s modulus of hPDMS was 4.7 MPa. Flat stamps for microcontact printing were made from Sylgard 184 (Dow Corning Inc). Fluorescently labeled (Alexa Fluor 488 NHS ester; Invitrogen) fibrinogen (Sigma) was physisorbed from solution (0.1 mg/ml in PBS) onto stamps for one hour, quickly washed with distilled water, and blown dry under nitrogen. Coated stamps were brought into conformal contact with micropost arrays, which had been pre-activated for 7 minutes by UV/Ozone treatment, pressed down using a forceps, and removed. The transfer efficiency of fibrinogen was confirmed by fluorescence microscopy of stamps and microposts and typically was >90%. Micropost arrays were passivated using a 1:2 mix (0.5 mg/ml in PBS) of fluorescently labeled (DyLight 405 NHS ester; Fisher Scientific Ireland) endotoxin-free bovine serum albumin (BSA; Sigma) to non-labeled BSA for 30 minutes, followed by Pluronics F127 (0.5% w/v in PBS) for 30 minutes, before washing three times with PBS. Washed platelets were resuspended in Tyrode’s buffer and seeded in the presence of thrombin (0.01 U/ml) onto the coverslips at a density of ca 2 million cm^-2^ for 60 minutes at 37°C, fixed in 3% paraformaldehyde in PBS for 15 minutes, and washed three times in PBS. Samples were permeabilized, blocked and stained for F-actin (phalloidin-Alexa Fluor 647; Invitrogen; 1:100 in 3% BSA in PBS) for 30 minutes, washed, and mounted in a chamber (Chamlide; Live Cell Instruments Co Ltd) containing PBS. Confocal images were obtained on a Leica SP8 from top and bottom slices of the microposts. Image analysis of traction forces was performed by custom MATLAB code. Posts were detected by template matching and their positions refined by a radial symmetry fit. Post deflections were deduced from the positions of the same post in top and bottom slices after removal of systematic offsets between the slices. Traction forces were calculated from the post deflections by Hooke’s law using a spring constant of 34.51 nN/μm (*38*).

### Scanning ion conductance microscopy (SICM)

35 mm round polystyrene cell culture dishes (Greiner Bio One, Ref. 627161) were coated with fibrinogen (Sigma-Aldrich, F3879, 0.1 mg/ml) by incubation for 1 h at 37°C. Washed mouse platelets were resuspended in Tyrode’s buffer supplemented with 1 mM Ca^2+^ and seeded in the presence of thrombin (Sigma-Aldrich, T6884, 0.01 U/ml) for 15 min at room temperature. Afterwards, the culture dishes were washed three times with Tyrode’s buffer to remove non-adherent platelets. The dishes were then installed in custom-built SICM setups (*39*) and imaged within one hour. Platelet morphology and elastic modulus were visualized using borosilicate pipettes with an inner radius of ≈90 nm and an applied pressure of 10 kPa. Imaging was done at a 20 Hz pixel rate, with 32 x 32 or 64 x 64 pixels at scan sizes between 6 x 6 and 12 x 12 μm^2^. Data were pooled from two donors. Analysis of elastic modulus *E* was calculated as described before (*40*).

### Thromboelastography

Rotational thromboelastometry (ROTEM) was used with a modified version of the commercial tissue factor-activated test (EXTEM) to detect changes in fibrinolytic activity (*41*).

In detail, we triggered the lysis *ex vivo* by adding rtPA (Abcam) at a final concentration of 137 ng/ml as reported (*41*). In a parallel application, we tried to contrast the lysis stimulation by contemporarily adding tranexamic acid (Merck) at a final concentration of 100 μM, which is expected to inhibit fibrinolysis (*31*). The ROTEM device was from Pentapharm GmbH, Munich, Germany. The maximum runtime was set to 150 minutes. The following standard parameters were analyzed: lysis onset time (LOT), and lysis time (LT) in minutes.

### Data analysis

Results are from at least two independent experiments per group, if not otherwise stated. Correction for multiple comparisons was analyzed using the Holm-Sidak method and differences between control and mutant sample were statistically analyzed using the Mann-Whitney-U test. P-values < 0.05 were considered as statistically significant: *0.05 > p ≥ 0.01; **0.01 > p ≥ 0.001; ***p < 0.001. Results with a P-value ≥ 0.05 were considered as not significant.

## Acknowledgments

The authors thank Daniela Naumann, Kim Ulbrich, Tamara Nahm, Doreen Biedenweg, Manu Emanuel, Nguyen Thi-Huong, Julia Klauke, Ricarda Raschke and Jan Wesche for excellent technical assistance.

## Funding

This work was supported by TR240 grant with project number 374031971 of the Deutsche Forschungsgemeinschaft (DFG; German Research Foundation), by the Deutsches Zentrum für Herz-Kreislauf-Forschung (Postdoc start-up grant under grant agreement 81X3400107) and by the Bundesministerium für Bildung und Forschung (ZIK grant under grant agreement 03Z22CN11). I.S. acknowledges funding from RCSI.

## Author contributions

Performed experiments: JB, LS, OO, IS, CZ, RK, HE, JR, ZN, RP, MB

Data analysis: all authors

Writing – original draft: JB, LS, RP, MB

Supervised research: OO, AG, RP, MB

Funding acquisition: OO, IS, TES, RP, MB

All authors have critically revised and approved the final version of the manuscript.

## Competing interests

O.O. is co-founder and shareholder of Zellmechanik Dresden distributing the AcCellerator for real-time fluorescence and deformability cytometry measurements. All other authors declare no competing financial interests.

## Data and materials availability

All data needed to evaluate the conclusions in the paper are present in the paper and/or the Supplementary Materials and raw data are available from the corresponding authors upon request.

## Notes

### Competing Interest Statement

Oliver Otto is co-founder and shareholder of Zellmechanik Dresden distributing the AcCellerator for real-time fluorescence and deformability cytometry measurements. All other authors declare no competing financial interests.

