## Supplemental Information for "Reduced platelet forces underlie impaired hemostasis in mouse models of *MYH9*-related disease"

[1]Institute of Experimental Biomedicine – Chair I, University Hospital and Rudolf Virchow Center, Würzburg, Germany. [2]Institute for Immunology and Transfusion Medicine, University Medicine Greifswald, Greifswald, Germany. [3]Zentrum für Innovationskompetenz – Humorale Immunreaktionen bei Kardiovaskulären Erkrankungen, University Greifswald, Greifswald, Germany. [4]School of Pharmacy and Biomolecular Sciences, Irish Centre for Vascular Biology, Royal College of Surgeons in Ireland, Dublin, Ireland. [5]University of Pavia, Pavia, Italy. [6]Institute of Applied Physics, University of Tübingen, Tübingen, Germany.

†These authors contributed equally to this work.

*Correspondence to:
Markus Bender:
Raghavendra Palankar:

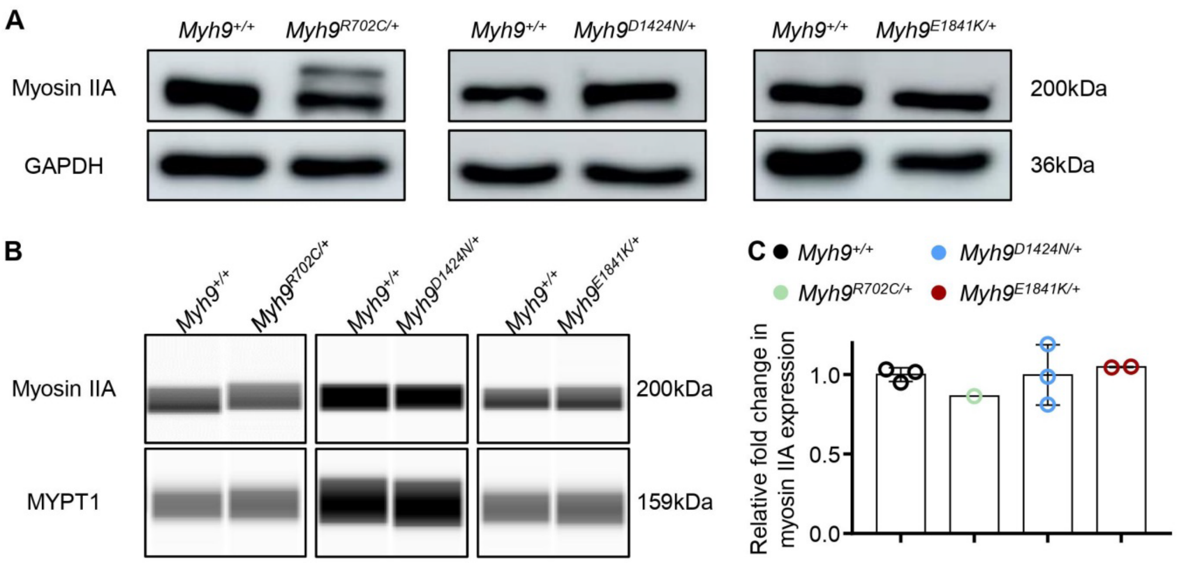


**Supplemental Fig. 1. Myosin IIA is expressed in platelets of *Myh9* mutant mice.** (**A**) Expression of myosin IIA in *Myh9*$^{D1424N/+}$, *Myh9*$^{E1841K/+}$ and corresponding control (*Myh9*$^{+/+}$) platelet lysates. Immunoblotting was performed on a 10% SDS-PAGE. Immunoblotting of *Myh9*$^{R702C/+}$ and control lysates was performed on a 7.5% SDS-PAGE. R702C mutant non-muscle myosin IIA-GFP and endogenous non-muscle myosin IIA could be detected. GAPDH served as loading control (n=3). (**B**) Myosin IIA and myosin phosphatase 1 (MYPT1) expression was confirmed using a capillary-based immunoassay approach. Fold change in myosin IIA expression relative to corresponding wildtype myosin IIA expression are shown in (**C**) for all mouse lines. Control shown corresponds to mouse line *Myh9*$^{D1424N/+}$. Each symbol represents one individual mouse.

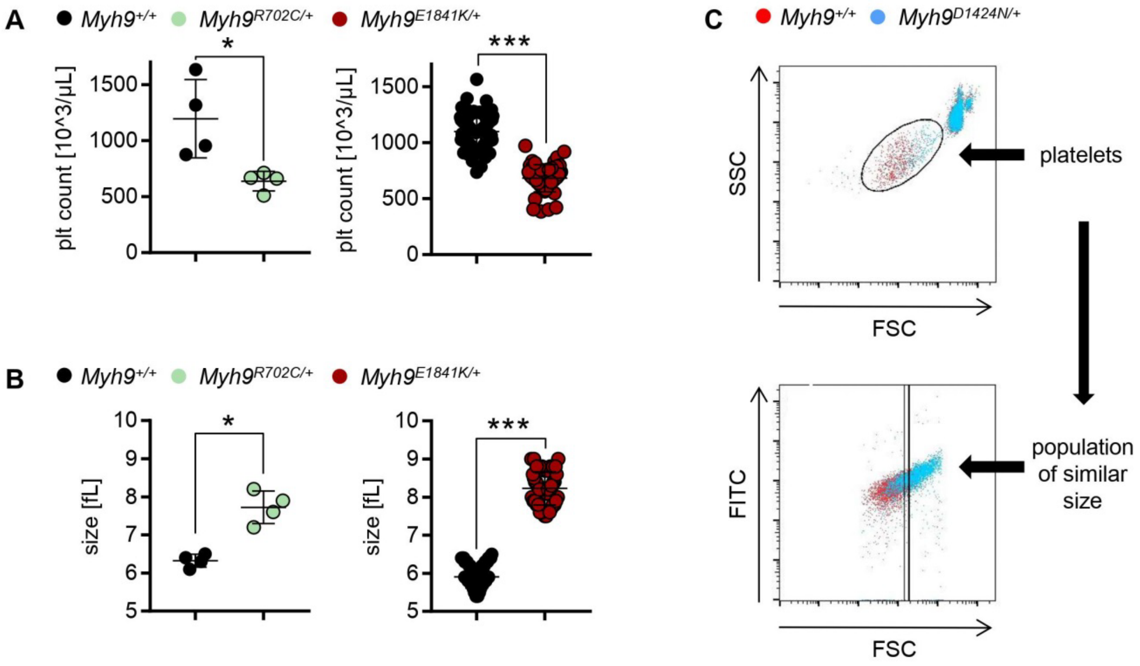


**D**

| | *Myh9*+/+ | *Myh9*R702C/+ | significance | *Myh9*+/+ | *Myh9*E1841K/+ | significance |
|---|---|---|---|---|---|---|
| GPIb | 10930 ± 461 | 12225 ± 737 | * | 8370 ± 296 | 9694 ± 746 | ** |
| GPV | 9337 ± 414 | 10129 ± 922 | ns | 5244 ± 1520 | 7442 ± 178 | ** |
| GPIX | 15084 ± 1491 | 15826 ± 1028 | ns | 7640 ± 4253 | 12564 ± 270 | * |
| αIIbβ3 | 10573 ± 500 | 10669 ± 320 | ns | 9882 ± 5074 | 13384 ± 759 | ns |
| α2 | 463 ± 24 | 483 ± 12 | ns | 1437 ± 93 | 1551 ± 76 | ns |
| CD9 | 27814 ± 775 | 29337 ± 1791 | ns | 32612 ± 2048 | 34219 ± 2280 | ns |
| GPVI | 1591 ± 213 | 1795 ± 174 | ns | 843 ± 479 | 1202 ± 74 | ns |
| CLEC-2 | 3158 ± 80 | 3498 ± 115 | *** | 1694 ± 1414 | 3583 ± 230 | * |

**Supplemental Fig. 2. Macrothrombocytopenia in R702C and E1841K mutant mice.** Determination of (**A**) platelet count and (**B**) platelet size. Each symbol represents one individual mouse (mean ± S.D.). (**C**) Gating strategy to determine platelet population of similar size for analysis of glycoprotein expression and platelet activation. (**D**) Glycoprotein expression on the platelet surface was determined by flow cytometry (n=4-7). Data are expressed as mean fluorescence intensity.

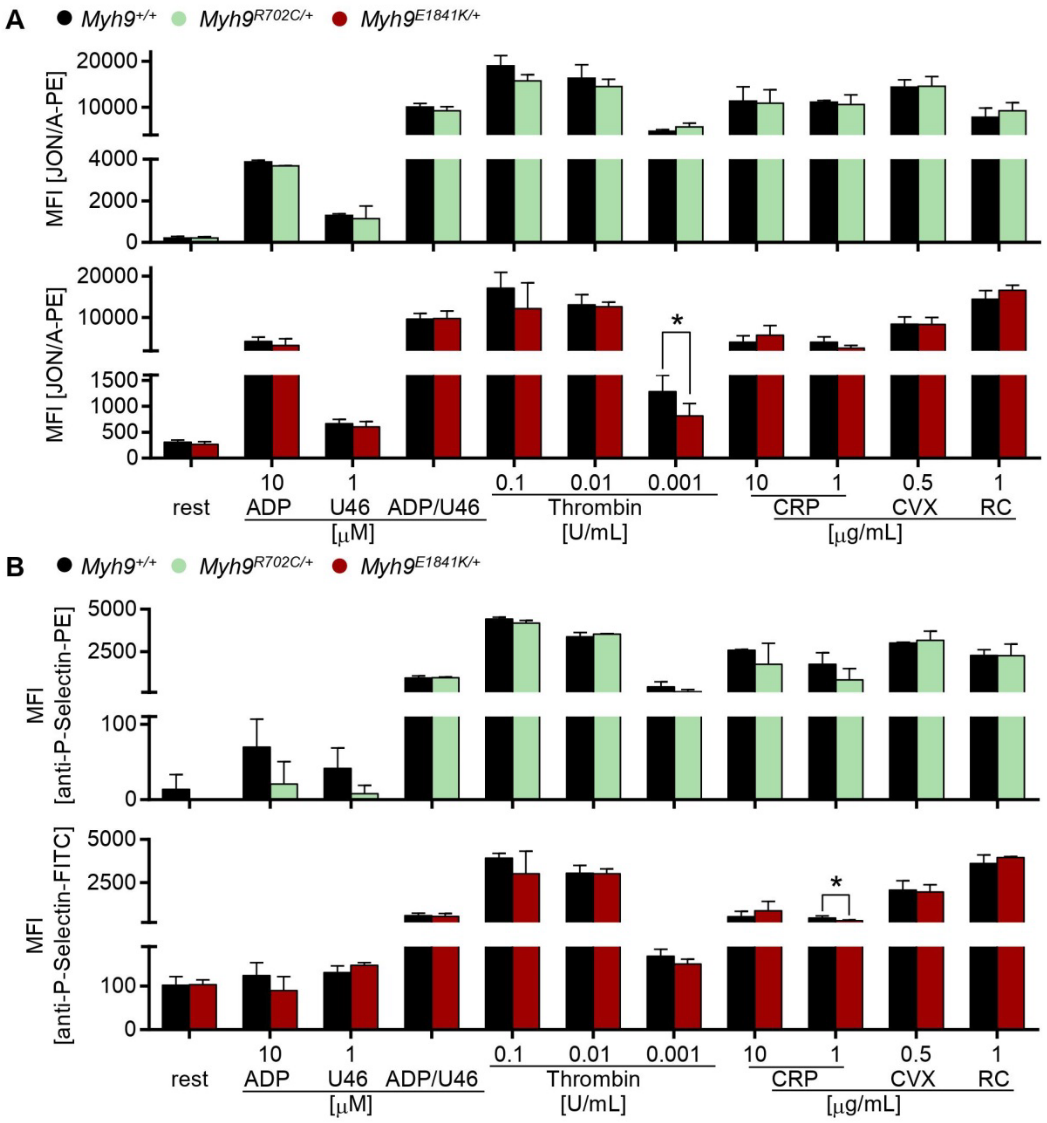


**Supplemental Fig. 3. Inside-out activation is unaltered in mutant platelets.** (**A**) Activation of platelet αIIbβ3-integrin (JON/A-phycoerythrin [PE]) and (**B**) α-granule release (anti-P-Selectin-fluorescein isothiocyanate [FITC]) under resting (rest) conditions and upon stimulation with different agonists (n=5, MFI). ADP: adenosine diphosphate; U46: thromboxane A2 analog U46619; CRP: collagen-related peptide; CVX: Convulxin; RC: Rhodocytin.

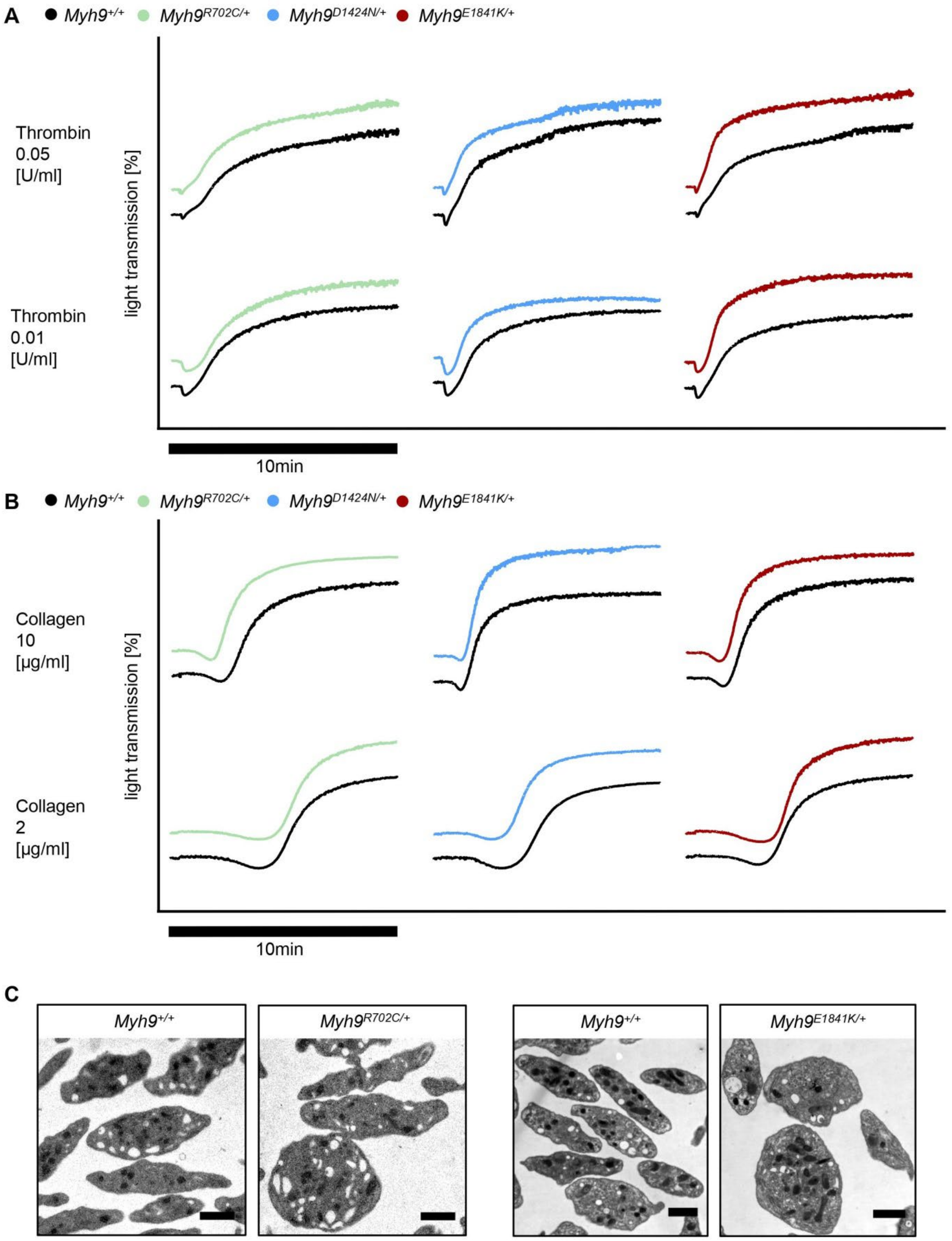


**Supplemental Fig. 4. Normal shape change and aggregation of *Myh9* mutant platelets.** Washed platelets were stimulated with (**A**) thrombin or (**B**) collagen and light transmission was recorded using a 4-channel aggregometer. Representative curves of two independent experiments are shown (n=2). (**C**) Representative transmission electron micrographs of mutant and control platelets. Scale bar: 1 µm.

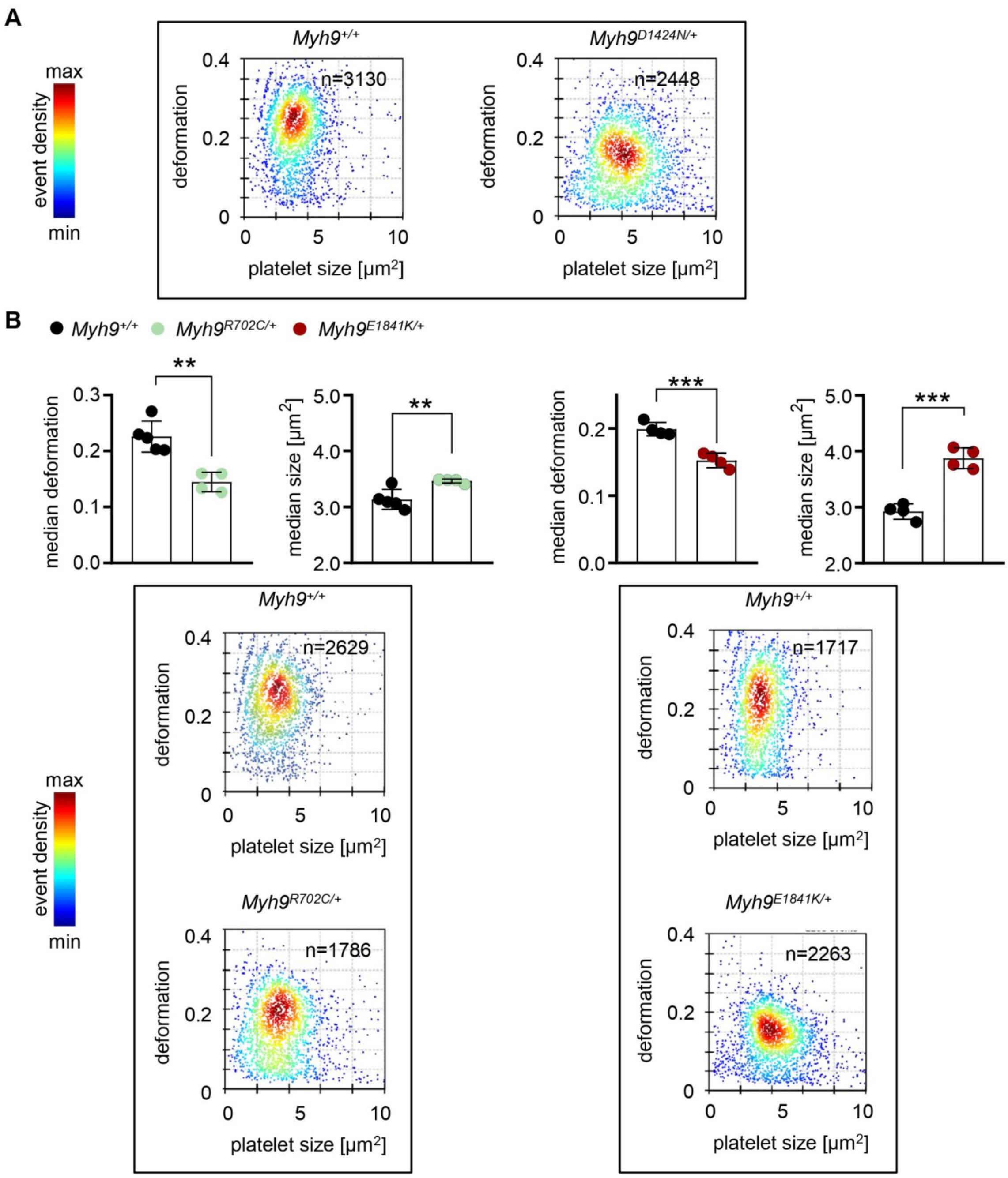


**Supplemental Fig. 5. Deformability measurements by RT-FDC.** (**A**) Representative KDE scatter plots from RT-FDC measurements displaying the distribution of single platelet deformability and their corresponding size between single platelets from *Myh9*$^{+/+}$ or *Myh9*$^{D1424N/+}$ mice (n= number of single platelets). (**B**) Each data point of RT-FDC measurement shows the median deformation and area of *Myh9*$^{+/+}$, *Myh9*$^{R702C/+}$ or *Myh9*$^{E1841K/+}$ platelets with at least 2000 platelets of one individual mouse. Bar plots show mean ± S.D. of platelet deformation or platelet area (n=4-5). Representative KDE scatter plots from RT-FDC measurements displaying the distribution of single platelet deformability and their corresponding size between single platelets from *Myh9*$^{+/+}$, *Myh9*$^{R702C/+}$, or *Myh9*$^{E1841K/+}$ mice (n= number of single platelets).

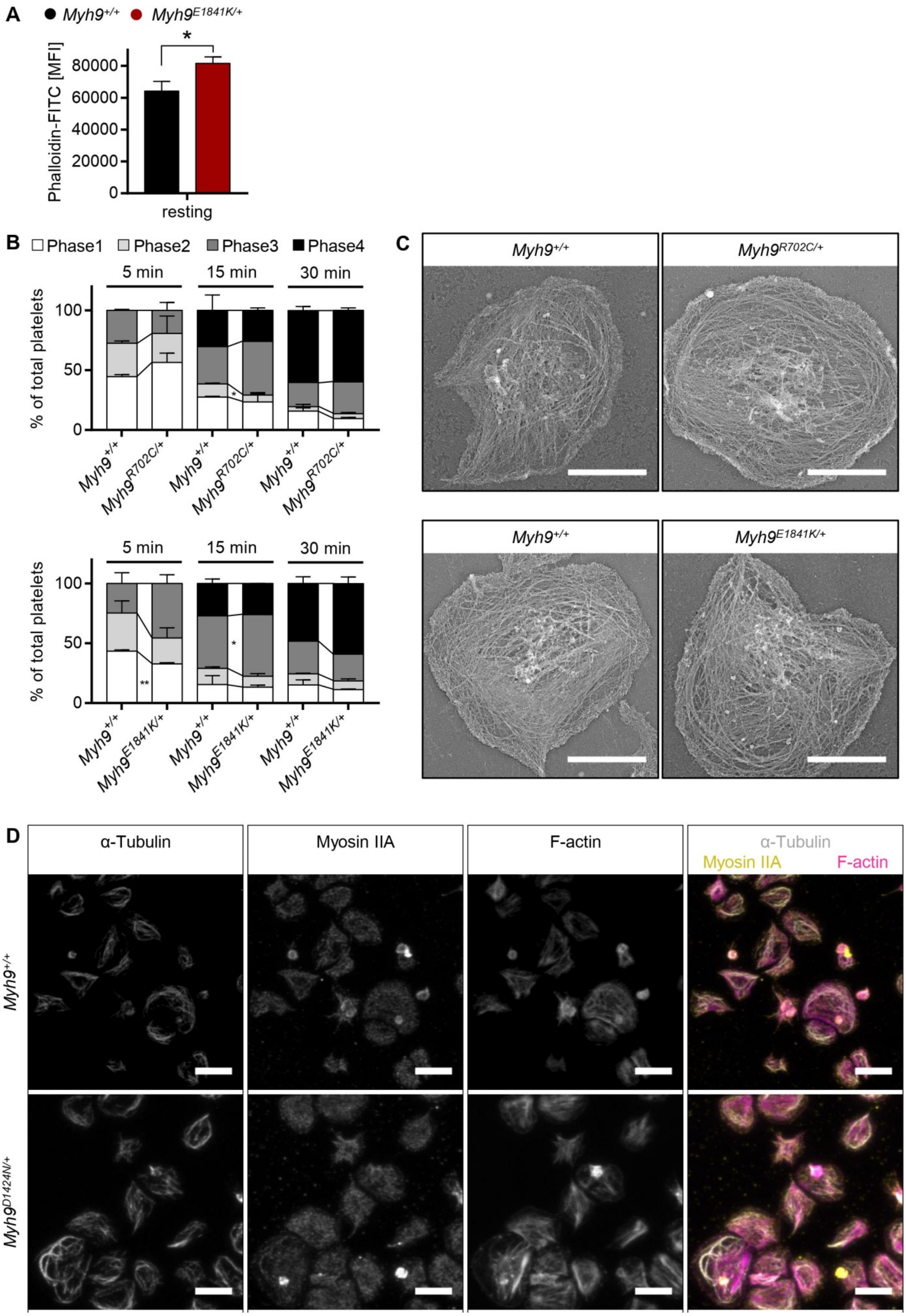


**Supplemental Fig. 6. Outside-in signaling of *Myh9* mutant platelets.** (**A**) F-actin content of resting platelets from mouse line E1841K was measured by flow cytometry after incubation with phalloidin-FITC (n=4). The mean fluorescence intensity is shown. (**B**) Platelets were spread on fibrinogen in the presence of thrombin and statistical analysis was performed of the different spreading phases of fixed platelets at different time points (n=2). Platelets of phase 1 were resting discoid shaped, phase 2 platelets formed filopodia, phase 3 platelets displayed lamellipodia and filopodia and phase 4 platelets were fully spread with only lamellipodia (mean ± S.D.). (**C**) Platinum replica electron microscopy of the cytoskeleton ultrastructure of spread platelets on fibrinogen was performed (scale bars: 2 µm). (**D**) Representative single stained and merged confocal images of *Myh9*$^{+/+}$ and *Myh9*$^{D1424N/+}$ platelets spread on fibrinogen in the presence of thrombin, stained for α-tubulin (grey), myosin IIA (yellow), and F-actin (magenta)(scale bars: 5 µm).

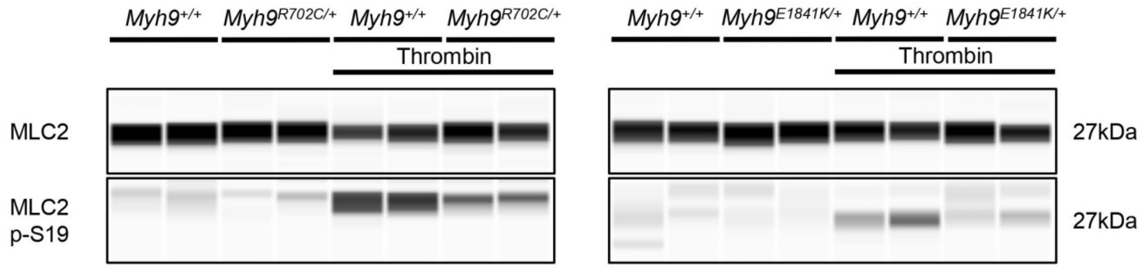


**Supplemental Fig. 7. Reduced myosin light chain 2 phosphorylation in *Myh9* mutant platelets.** Expression of myosin light chain 2 (MLC2) and phosphorylated myosin light chain 2 (MLC2p-S19) of resting and thrombin-activated (0.05 U/mL, 1 min) platelets was determined. Representative immunoblot of two independent experiments are shown (n=2).

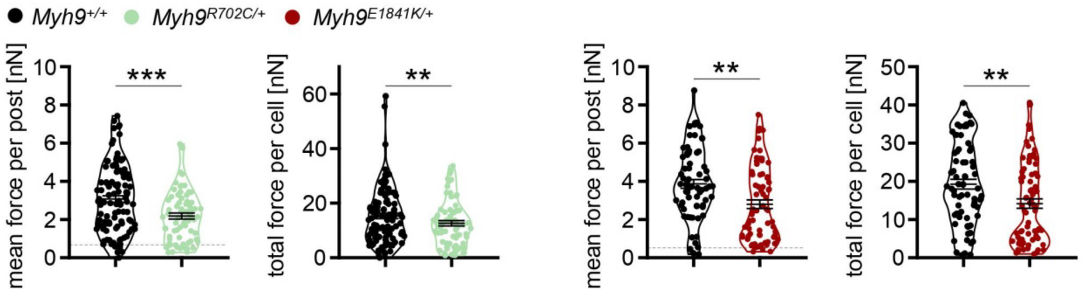


**Supplemental Fig. 8. *Myh9* mutant platelets show less contractile forces on micropost arrays.** Platelet contractile forces were measured per post and the sum of posts per cell (mean ± S.D.; *Myh9*$^{+/+}$: n=106; *Myh9*$^{R702C/+}$*:* n=73; *Myh9*$^{+/+}$: n=69; *Myh9*$^{E1841K/+}$*:* n=74).

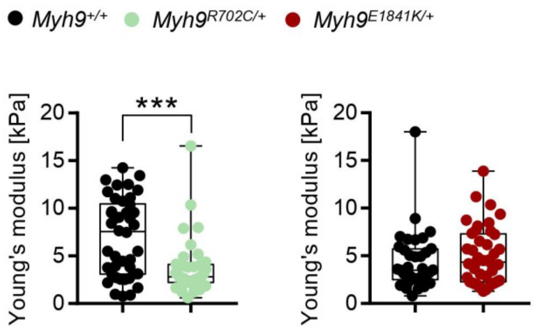


**Supplemental Fig. 9. Spread platelets of *Myh9*$^{R702C/+}$ but not *Myh9*$^{E1841K/+}$ mice are softer.** Washed *Myh9*$^{+/+}$, *Myh9*$^{R702C/+}$ and *Myh9*$^{E1841K/+}$ platelets spread on fibrinogen in the presence of thrombin were analyzed of their Young´s modulus using scanning ion conductance microscopy. Each symbol represents one platelet (*Myh9*$^{+/+}$: n=42 platelets; *Myh9*$^{R702C/+}$*:* n=36 platelets; *Myh9*$^{+/+}$: n=36 platelets; *Myh9*$^{E1841K/+}$*:* n=41 platelets) and box plots represents median ± S.D.

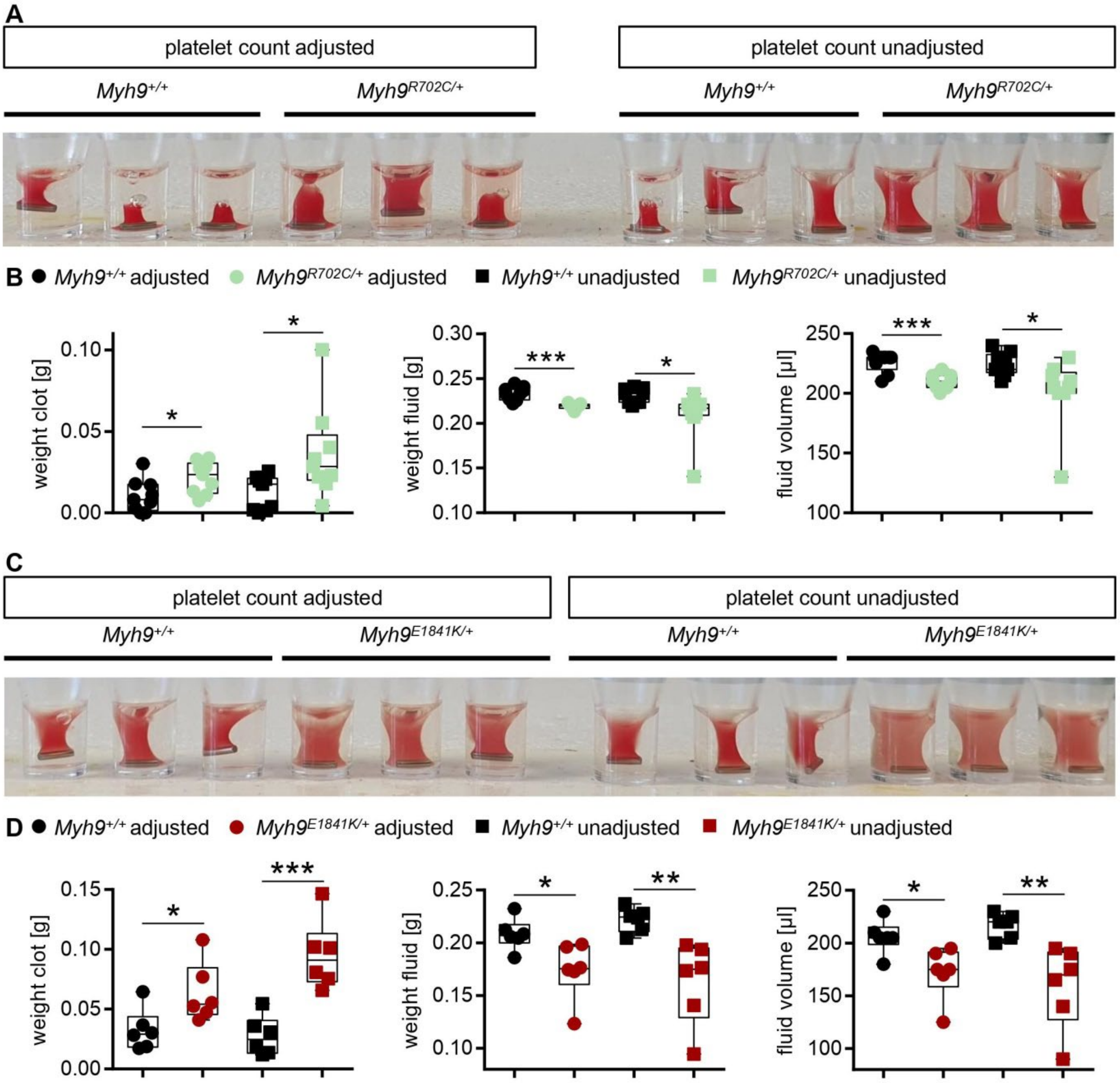


**Supplemental Fig. 10. Reduced extent of clot retraction in *Myh9* mutant samples.** (**A**) Representative image of clot formation of *Myh9*$^{+/+}$ and *Myh9*$^{R702C/+}$ samples at time point 60 minutes (n=3). (**B**) Results are median ± S.D. Each symbol represents one individual mouse. (**C**) Representative image of clot formation of *Myh9*$^{+/+}$ and *Myh9*$^{E1841K/+}$ samples at time point 60 minutes (n=3). (**D**) Results are median ± S.D. Each symbol represents one individual mouse.

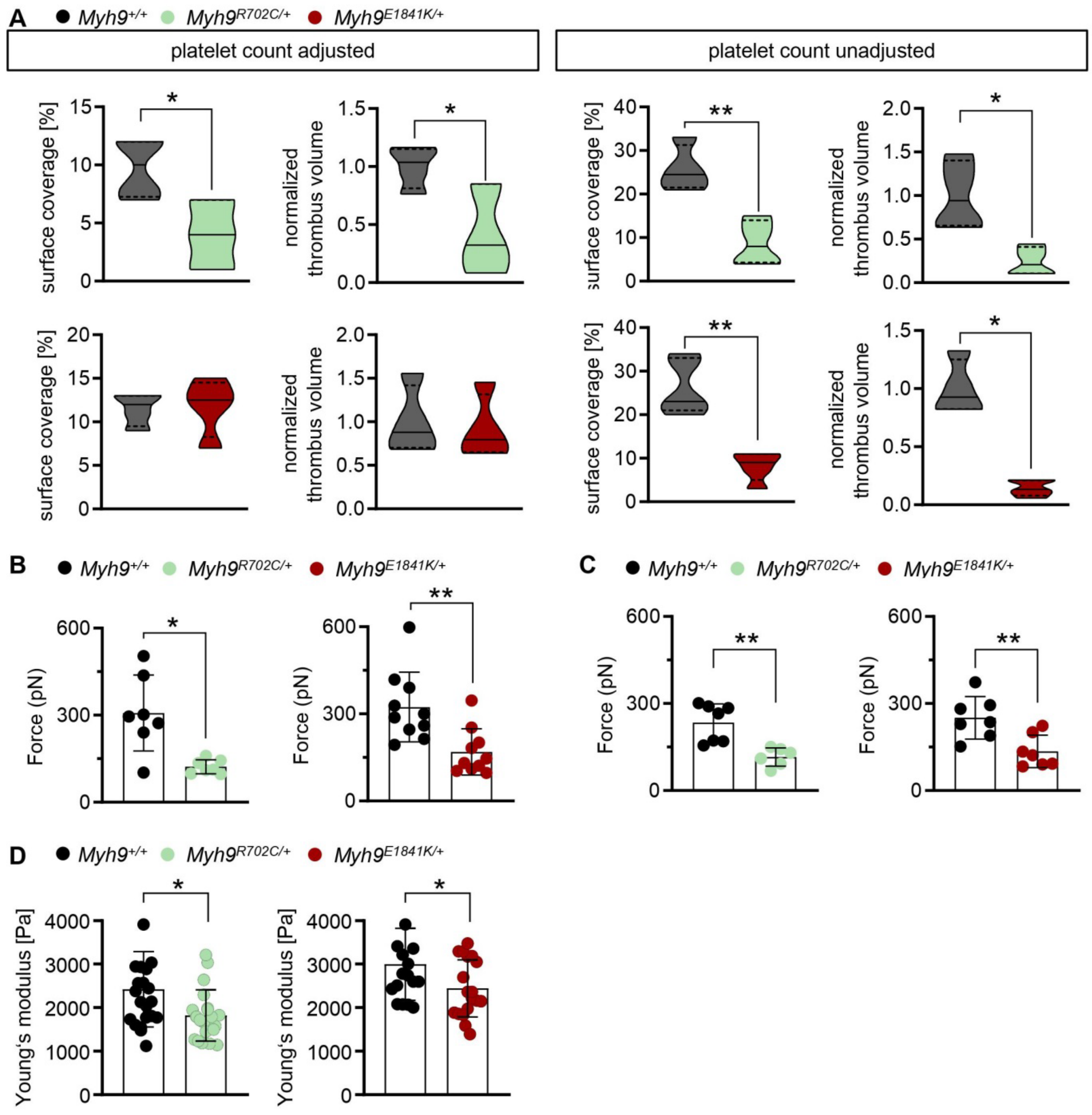


**Supplemental Fig. 11. Reduced thrombus formation under flow of *Myh9* mutant platelets.** (**A**) Assessment of platelet adhesion and aggregate formation under flow (1000/s) on collagen of *Myh9*$^{+/+}$ and *Myh9*$^{R702C/+}$ samples (upper row), and of *Myh9*$^{+/+}$ and *Myh9*$^{E1841K/+}$ samples (lower row). Analysis of the surface area covered by platelets (%) and the relative normalized thrombus volume are shown for platelet count adjusted conditions on the left side (n=4, mean ± S.D.) and unadjusted conditions on the right side (n=4-5, mean ± S.D.). Single platelet force spectroscopy of (**B**) platelet adhesion to collagen and (**C**) platelet-platelet interaction forces. (**B** to **C**) Each data point of summary graphs (mean ± S.D.) shows one platelet to collagen (n=7-10) or platelet to platelet (n=7) interaction. (**D**) Each data point of colloidal probe spectroscopy shows the median Young´s modulus of one *Myh9*$^{+/+}$, *Myh9*$^{R702C/+}$ or *Myh9*$^{E1841K/+}$ aggregate and bar plots show mean ± S.D. of Young´s modulus (n=4-5).

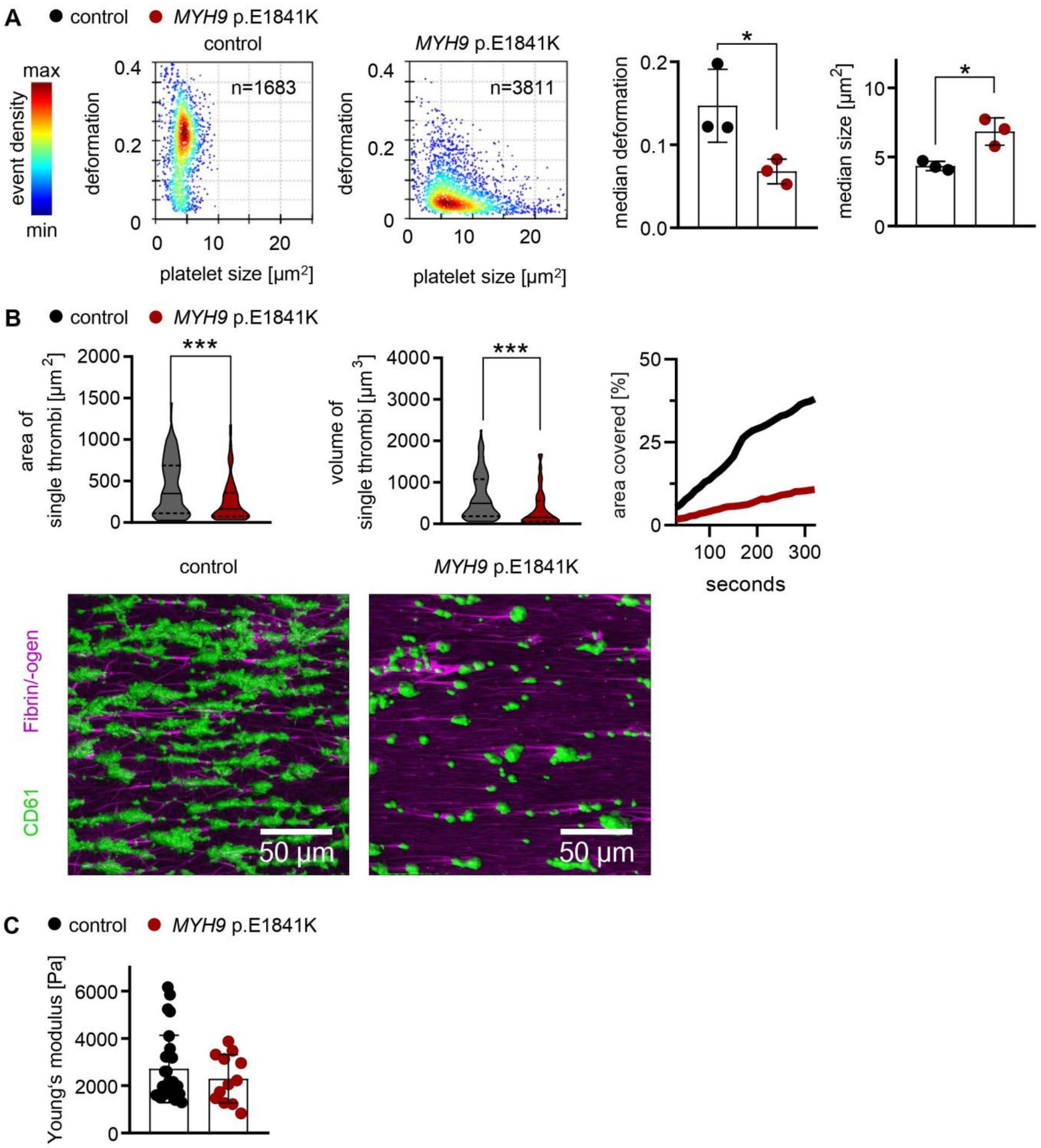


**Supplemental Fig. 12. *MYH9* p.E1841K patient platelets form smaller thrombi under flow.** (**A**) Representative KDE scatter plots from RT-FDC measurements displaying the distribution of single platelet deformability and their corresponding size between single platelets from a healthy individual (control) and from a *MYH9* p.E1841K patient (n= number of single platelets). Summary data points show the median values of individual donors and one *MYH9* p.E1841K patient on three different days and bar plots show mean ± S.D. of platelet deformation and size from non-stimulated platelets. (**B**) Platelet adhesion and aggregate formation under flow (1000/s) on collagen of human patient samples were assessed by a flow chamber assay. Area and volume of single thrombi are shown are shown as median ± quartiles of platelet count unadjusted conditions (n=67 single thrombi). Area covered over time shows the mean of thrombi from control and patient blood. Representative images taken at the end of the perfusion time (20 minutes) are shown in fluorescent images with platelets labeled with the anti-CD61-antibody and labeled fibrin/-ogen (scale bars 50 µm). (**C**) Whole blood from healthy individuals and *MYH9* p.E1841K patient was examined by colloidal probe spectroscopy. Each data point shows the median Young´s modulus of one aggregate and bar plots show mean ± S.D. of Young´s modulus.

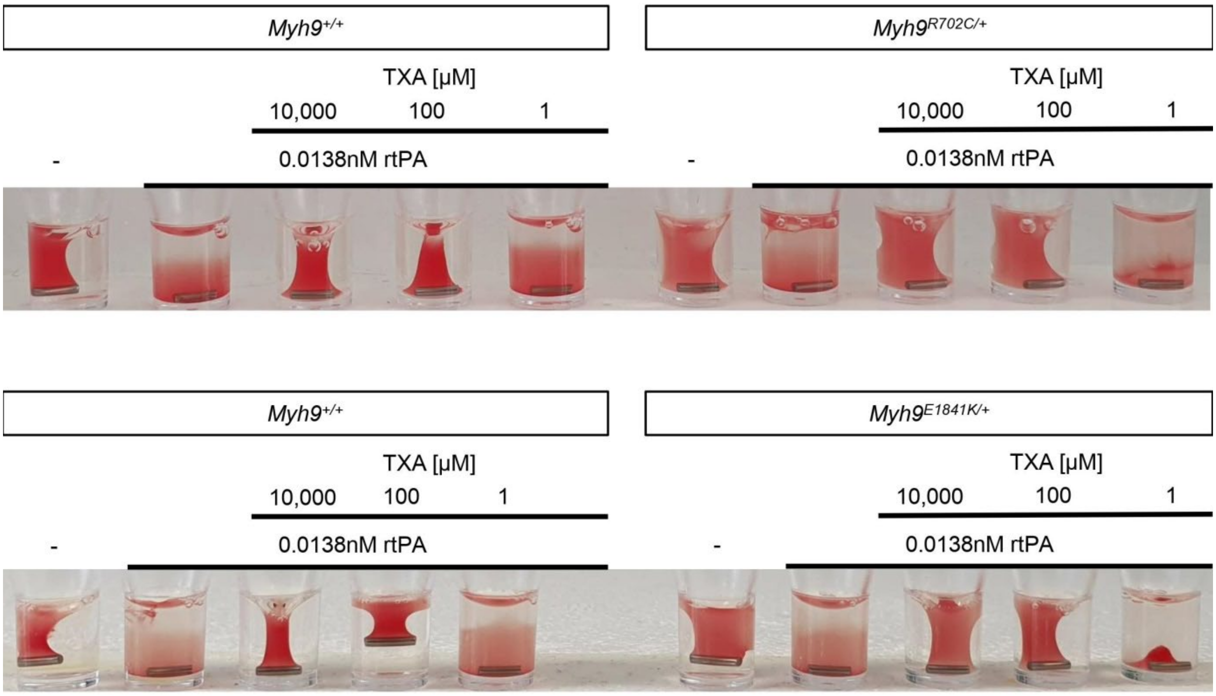


**Supplemental Fig. 13. Clot lysis induced by rtPA can be reversed by high concentrations of TXA.** Representative image (three independent experiments) of clot retraction of *Myh9*$^{+/+}$, *Myh9*$^{R702C/+}$, and *Myh9*$^{E1841K/+}$ samples treated with rtPA in a threshold concentration where lysis occurs. Addition of TXA at different concentrations.

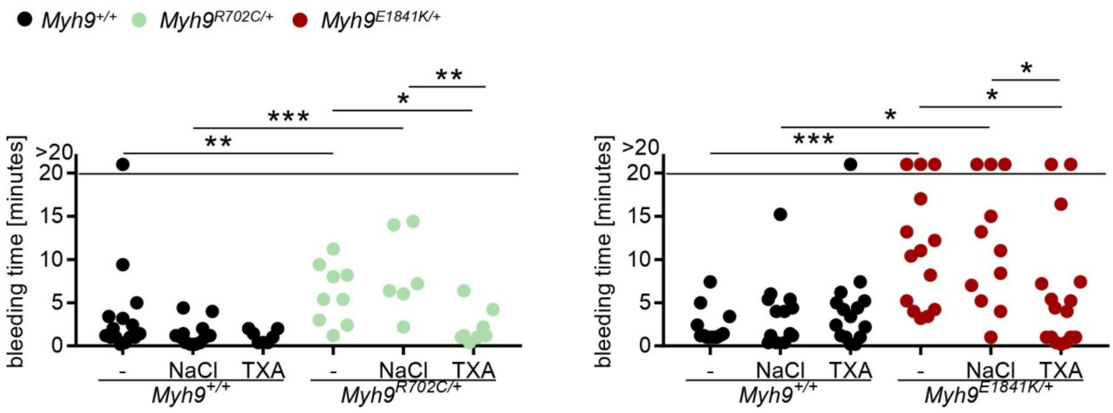


**Supplemental Fig. 14. Tranexamic acid improves hemostasis.** Tail bleeding times on filter paper of *Myh9* mutant mouse lines R702C and E1841K. Injection of sodium chloride served as control. Each symbol represents one individual mouse (mean ± S.D.). Mann-Whitney-U test was used to analyze data excluding data points exceeding 20 minutes.

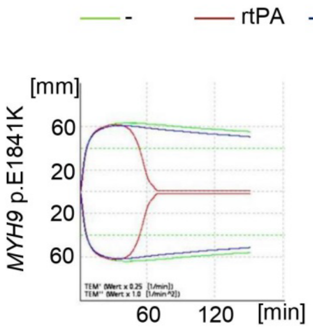


| | LOT<br>- | LOT<br>rtPA | LOT<br>rtPA+TXA |
|---|---|---|---|
| *MYH9* p.E1841K | >150‘ | 46‘13“ | 136‘27“ |

| | LT<br>- | LT<br>rtPA | LT<br>rtPA+TXA |
|---|---|---|---|
| *MYH9* p.E1841K | >150‘ | 63‘ | >150‘ |

**Supplemental Fig. 15. Shortened LOT and LT of *MYH9* p.E1841K sample can be reverted by TXA.** Lysis onset time (LOT) and lysis time (LT) of human sample assessed with ROTEM. Overlapping of the modified ROTEM analysis curves (green: no treatment; red: rtPA stimulation; blue: rtPA and TXA stimulation).

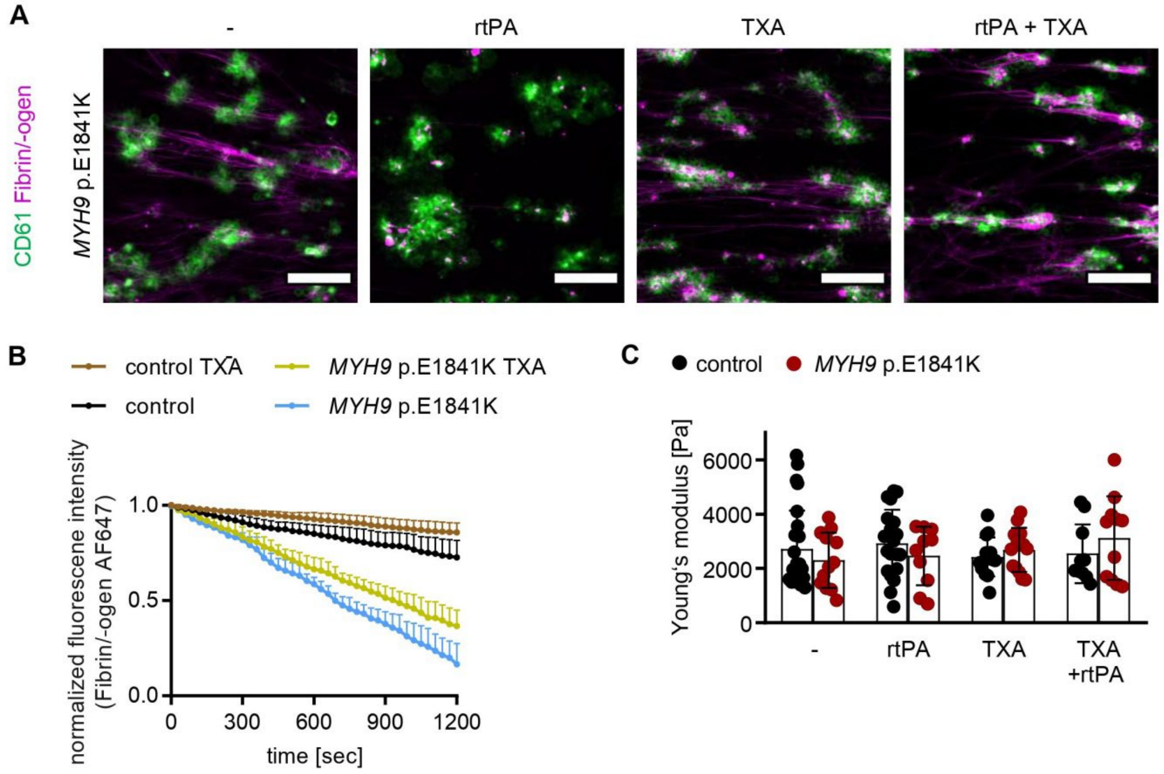


**Supplemental Fig. 16. Fibrinolysis in a microfluidic flow chamber decreased by TXA.** (**A**) Representative fluorescence images of human platelet thrombi (CD61; green) and fibrin (magenta) at 20 minutes after in microfluidic flow chamber at a shear of 100 $s^{-1}$ (scale bars 50µm) and (**B**) Time course of changes in stability of fibrin/-ogen (normalized fluorescence of fibrin/-ogen AF647, mean ± S.D.) on sites of platelet thrombi after addition of TXA (100 µM)

compared to non-treated control. (**C**) Thrombus stiffness in whole blood from healthy individual and *MYH9* p.E1841K patient was examined (untreated, with rtPA, with TXA and with TXA + rtPA) using colloidal probe spectroscopy. Each data point shows the median Young´s modulus of one healthy individual or *MYH9* p.E1841K aggregate and bar plots show mean ± S.D. of Young´s modulus.
